# Sex-biased Transcriptome in *in vitro* Produced Bovine Early Embryos

**DOI:** 10.1101/2025.03.04.641446

**Authors:** Meihong Shi, Guangsheng Li, Hannah Marie Araujo, Angie S Lee, Jingzhi Zhang, Yoke Lee Lee, IISAGE Consortium, Soon Hon Cheong, Jingyue (Ellie) Duan

## Abstract

**Background:** Morphologic sex differences between males and females typically emerge after the primordial germ cell migration and gonad formation, although sex is determined at fertilization based on chromosome composition. A key debated sexual difference is the embryonic developmental rate, with *in vitro* produced male embryos often developing faster. However, the molecular mechanisms driving early embryonic sex differences remain unclear.

**Results:** To investigate the transcriptional sex difference during early development, *in vitro* produced bovine blastocysts were collected and sexed by PCR. A significant male-biased development was observed in expanded blastocysts. Ultra-low input RNA-seq analysis identified 837 DEGs, with 231 upregulated and 606 downregulated in males. Functional enrichment analysis revealed male-biased DEGs were associated with metabolic regulation, whereas female-biased DEGs were related to female gonad development, sex differentiation, inflammatory pathways, and TGF-beta signaling. Comparing X chromosome and autosome expression ratio, we found that female-biased DEGs contributed to the higher X-linked gene dosage, a phenomenon not observed in male embryos. Moreover, we identified the sex-biased transcription factors and RNA-bind proteins, including pluripotent factors such as *SOX21* and *PRDM14*, and splicing factors *FMR1* and *HNRNPH2*. Additionally, we revealed 1,555 significantly sex-biased differential alternative splicing (AS), predominantly skipped exons, mapped to 906 genes, with 59 overlapping with DEGs enriched in metabolic and autophagy pathways. By incorporating novel isoforms from long reads sequencing, we identified 1,151 sex-biased differentially expressed isoforms (DEIs) associated with 1,017 genes. Functional analysis showed that female-biased DEIs were involved in the negative regulation of transcriptional activity, while male-biased DEIs were related to energy metabolism. Furthermore, we identified sex-biased differential exon usage in *DENND1B, DIS3L2, DOCK11, IL1RAPL2,* and *ZRSR2Y,* indicating their sex-specific regulation in early embryo development.

**Conclusion:** This study provided a comprehensive analysis of transcriptome differences between male and female bovine blastocysts, integrating sex-biased gene expression, alternative splicing, and isoform dynamics. Our findings indicate that enriched metabolism processes in male embryos may contribute to the faster developmental pace, providing insights into sex-specific regulatory mechanisms during early embryogenesis.

**Plain English summary:** Male and female early embryos develop at different speeds, with male embryos often developing faster than female embryos. However, the reasons behind these early differences remain unclear. In this study, we examined gene activity in bovine embryos to uncover the biological factors regulating these early sex differences. We collected in vitro-produced bovine blastocysts, examined their sex, and confirmed that male embryos develop faster. By analyzing global gene activity, including alternative splicing, which allows one gene to code for multiple RNA isoforms and proteins, we found distinct gene expression profiles between male and female embryos. Male embryos showed higher activity in genes related to metabolism and cellular functions, while female embryos had increased activity in genes associated with female-specific gonad development and gene expression regulation. We also examined differences in how genes on the X chromosome were expressed. Female embryos had higher X-linked gene expression, which may contribute to sex-specific developmental regulation. Additionally, we identified sex-specific transcription factors and RNA-binding proteins that regulate early embryo development, some of which are known to control pluripotency and gene splicing. Overall, our study provides new insights into how gene activity shapes early sex differences, suggesting that enhanced metabolism in male embryos may be a key driver of their faster developmental rate.

**Highlights:**

- Male embryos develop faster due to increased gene expression in metabolism pathways
- Female embryos exhibit higher X-linked gene expression, suggesting X-dosage compensation plays a role in early development
- Sex-biased alternative splicing events contribute to embryonic metabolism, autophagy, and transcriptional regulation in embryos
- Sex-biased isoform diversity contributes to distinct developmental regulation in male and female embryos
- Key pluripotency factors (SOX21, PRDM14) and splicing regulators (FMR1, HNRNPH2) drive sex-specific gene expression

## Background

Assisted reproductive technologies (ARTs), particularly *in vitro* fertilization (IVF), have been widely used to improve livestock fertility, herd genetics, and production. One interesting observation in these technologies is the differing developmental pace between male and female embryos during the early stages of embryogenesis, even before gonad differentiation occurs [1, 2]. This difference has been observed in various species over decades, including humans [3–9], mice [10–12], cattle [1, 13, 14], pigs [15, 16], and sheep [17]. However, the findings are controversial, as some studies report no significant sex biases in blastocyst formation and birth count, particularly in human IVF embryos [18–20]. Culture conditions and experimental design, such as differences in IVF methods, culture medium supplements, and *in vivo* or *in vitro* developmental environment, may contribute to these discrepancies [21, 22].

Previous studies have identified some sex-specific differences in early embryos prior to gonadal differentiation, such as differential expression of imprinted and sex-linked genes, glucose metabolism related activities, mitochondrial DNA (mtDNA) copy numbers, and telomere lengths [23–28]. Recent advances in transcriptomics, including microarray and RNA sequencing (RNA-seq) using embryos fertilized by sex-sorted semen or in single embryos, have provided deeper insights into these sex-specific differences at the transcriptomic level [29–33]. For example, *in vivo* produced bovine male embryos show a different transcriptome profile [30, 31], having a higher expression of pluripotent factors and a lower expression of genes related to glucose transport and apoptosis compared to females [34]. Besides, single-cell RNA-seq in human embryos indicated a distinct transcriptomic dynamic between female and male embryos after embryonic genome activation [32]. Additionally, the pig embryos also showed distinct transcriptional differences between sexes with a dynamic compensation of X chromosome in the female embryos [33]. Despite these findings, the underlying molecular mechanisms driving sex-specific differences in early embryogenesis remain poorly understood. Moreover, the use of embryos produced with sex-sorted semen introduces additional variability [34], as the presence of mixed sexed embryos can influence results. Thus, a comprehensive investigation of transcriptomic differences between *in vitro* produced male and female early embryos with precisely defined sex is essential to address the knowledge gap in sex-specific embryogenesis.

Regulation of transcription is mediated by multiple mechanisms, including transcription factors (TFs), TF-associated cofactors, and alternative splicing (AS). TFs are DNA-binding proteins that activate or suppress transcription, playing critical roles in gene regulation across evolution, development, and diseases [35]. Alternative splicing, controlled by spliceosome complex, allows a single gene to generate multiple transcript isoforms [36], significantly expanding transcriptomic diversity. Understanding the interplay between sex-specific gene expression regulation, alternative splicing events, and isoform diversities in early embryos is critical to unraveling the molecular basis of sex-specific development.

Beyond autosomal regulations, the regulation of genes located on sex chromosomes also has significant contributions to sex determination and sex-specific development. In mammals, the XY sex-determination system requires a balance of gene dosage between males (XY) and females (XX), and between sex chromosomes and autosomes [37]. Mechanisms such as X-chromosome inactivation (XCI) in females and X-chromosome upregulation (XCU) in both sexes are essential to maintain this balance and prevent gene dysregulation [38]. In early mammalian embryogenesis, the exact timing and the extent of these processes remain unclear. While Xist expression begins early in development, partial XCI has been observed [39], with some of the X-linked genes escaping the XCI silencing [40, 41]. Whether X dosage compensation contributes to the sex-specific embryo developmental differences in bovine embryos remains unclear.

In this study, we aim to uncover genomic differences between male and female bovine embryos produced *in vitro*. Using PCR-based single embryo sex determination and comprehensive RNA-seq analysis, we identified the differential gene expression (DEG) and associated regulators, examined the X-chromosome dosage compensation, characterized the sex-biased alternative splicing events, and analyzed isoform diversities between male and female blastocysts. Our findings reveal that male embryos developed faster, driven by upregulated metabolism activities compared to female embryos, while female embryos were more active in immune response, female-specific gonadal development, and protein processing.

## Materials and methods

### Embryo Production and sample collection

Bovine embryos were produced from slaughterhouse ovaries using commercial media (IVF Bioscience, Falmouth, UK) according to the manufacturer protocols. Bovine ovaries were obtained from a slaughterhouse (Cargill, Wyalusing, PA) and transported to the lab in warm saline solution (0.9% Sodium chloride, Sigma-Aldrich) maintained between 23.5-30°C within 2 h. Follicles (diameter: 2 - 8 mm) were aspirated using an 18-gauge hypodermic needle attached to an aspiration vacuum pump. Cumulus-oocyte complexes (COCs) with homogenous cytoplasm and surrounded by at least three layers of cumulus cells were selected and matured in BO-IVM™ medium (IVF Bioscience, Falmouth, UK) for 22h at 38.5°C in a humidified atmosphere with 5% CO_2_.

Matured COCs were transferred to BO-IVF™ medium (IVF Bioscience, Falmouth, UK) and cocultured with thawed semen at a final concentration of 1× 10^6^ sperm/mL at 38.5°C in a humidified atmosphere with 5% CO_2_. Fertilization was performed with cryopreserved semen from bulls of known fertility generously donated by Genex Cooperative. Semen preparation was conducted using BO-SemenPrep™ medium and recovered sperm concentration was determined with a sperm cell counter (Nucleocounter SP-100, ChemoMetec, Allerod, Denmark) then used for fertilization. To improve genetic diversity, two batches of semen from two different bulls were rotated every week, and there was no difference in developmental rates between them. After 18 h of fertilization, presumptive zygotes were denuded by vortexing at 3,200 rpm for 60s twice in a wash medium (IVF Bioscience, Falmouth, UK). Zygotes with intact membranes were transferred into BO-IVC™ medium (IVF Bioscience, Falmouth, UK) and cultured in a humidified atmosphere (5% O_2_, 5% CO_2_, and 90%N_2_) at 38.5°C for further blastocyst collection.

On Days 7 and 8, non-expanded, expanded, and hatched blastocysts were washed three times with DPBS (Life Technology, USA), collected individually in PCR tubes, and snap-frozen at −80°C for later sex determination. Expanded blastocysts were placed individually in 10ul drops of DPBS covered with mineral oil in 35mm culture dishes after a DPBS wash and evenly divided into two halves with a microblade under the microscope. Each half contained both the inner cell mass and trophectoderm (**Figure 1A**). Cut embryos were stored separately at −80°C for future use.

**Figure 1.**
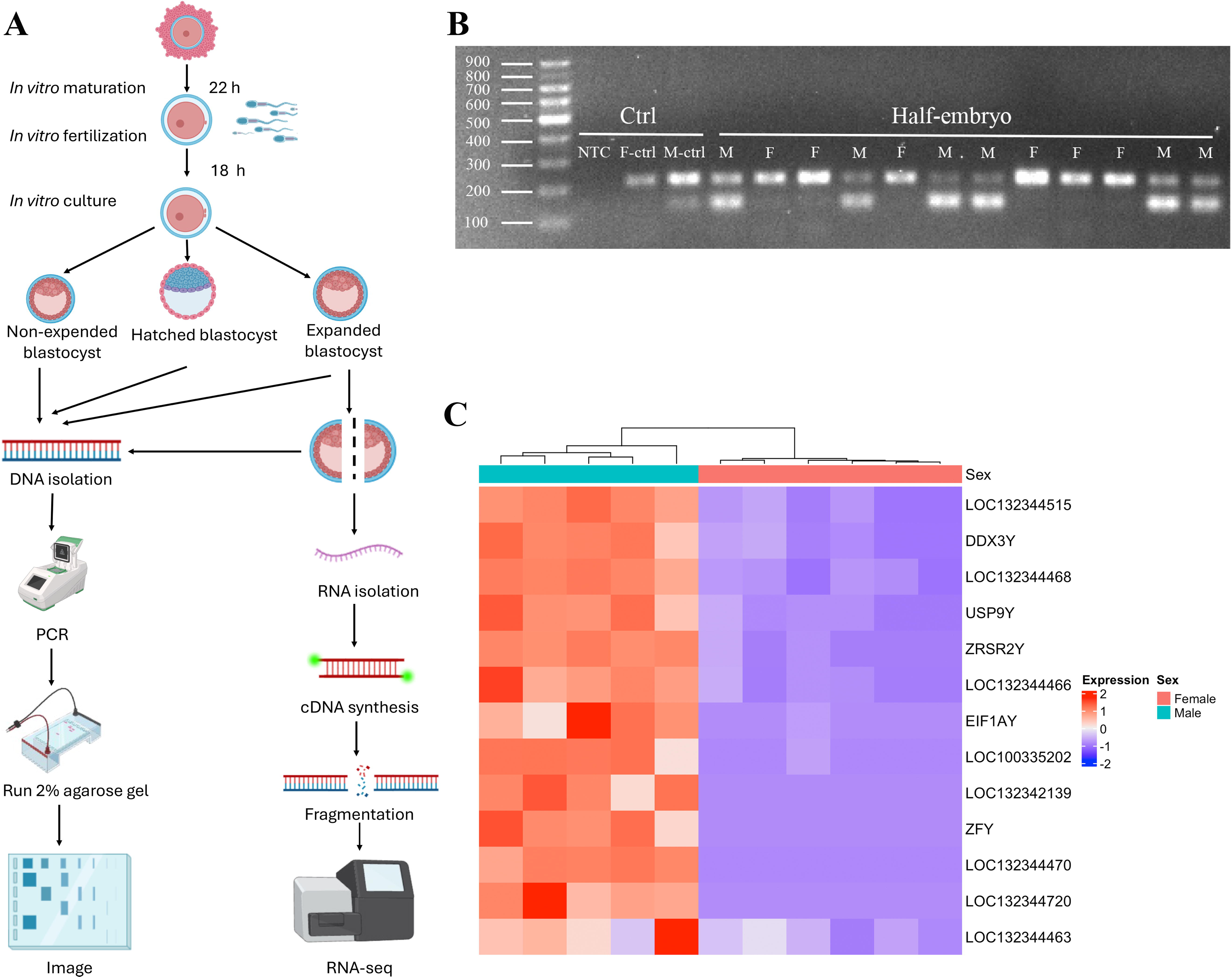
Overview of embryo sex determination. A) Schematic overview of the experimental setup. B) Agarose gel of amplified gDNA of half blastocysts. Each lane is one control sample or a half-cut embryo; NTC: no-template control; F-ctrl: gDNA from bovine mammary alveolar cells (MAC-T cells); M-ctrl: gDNA from bovine testis tissues. Identification of the autosomal amplicon (217 bp) located above and the Y-specific amplicon (143 bp) located below. C) Heatmap of all male-specific DEGs located on Y-chromosome in all RNA-seq samples.

### Sex determination

Embryo sexing followed a modified protocol [42]. Briefly, dissected embryos were lysed with 5 uL of cell lysis buffer with Proteinase K (1: 50 dilution, SingleShot Cell Lysis kit, 1725081, Bio-rad) and 100 ug/mL RNase (T3018L, New English Biolab), incubated at room temperature (RT) for 10 to 20 min, followed by 5 min at 37°C and 5 min at 75 °C.

For PCR, 12.5 uL of One*Taq*® Quick-Load® 2× Master Mix (M0486L, New England Biolabs) was mixed with 0.4 ul of 10 μM primers and 7.1 ul of nuclease-free water. The first round of PCR amplified a Y-linked DNA region using following conditions: 95°C for 5 min; 20 cycles of 95°C for 15 s, 58°C for 15 s, and 72°C for 15 s; then 10 min at 72°C. For the second round, 1 μL of 4 μM autosomal primers were added, and an additional 17 PCR cycles were performed under the same conditions. Primer sequences were listed in **Table S1.**

PCR products (5 μL) were mixed with 1 μL of 6× DNA loading dye (R0611, Thermo Scienticic) and loaded on 2% (wt/vol) agarose gels stained with SYBR Safe DNA gel Stain (S33102, Thermo Scienticic) for electrophoresis. Embryos were classified as males if the Y-linked and autosomal amplicons were both present and as females if only the autosomal amplicon was detected.

### RNA-seq library construction

Two embryos of the same sex, collected on the same day (days 7 or 8) were pooled. Five male and six female embryos were used for RNA-seq library construction. Libraries were constructed using SMART-Seq® HT PLUS Kit (R400748, Taraka). Briefly, pooled embryos were lysed in 1× Lysis buffer with RNase inhibitor, and full-length double-strand cDNA was synthesized and amplified with 15 cycles with a one-step master buffer. The cDNA was quantified by Fragment Analyzer 5200 (Aligent) and purified with AMPure XP beads (Beckman Coulter). For each replicate, 8 ng of cDNA was fragmented, stem-loop adapter was added, and followed by 13 cycles of PCR for library amplification with unique dual indexes. Libraries were quality controlled (QC) and sequenced at Novogene Inc. (Sacramento, CA, USA), using NovoSeq PE150 platform.

### RNA-seq analysis

The adaptor removal and QC of the raw sequencing reads were performed using fastp (v 0.20.0) [43]. Reads with a percentage of low-quality base (quality score <20) >40% were removed. Reads with length < 30 bp or with high N content (>5%) were also removed in this study. The cow reference genome (ARS-UCD 2.0) was downloaded from NCBI database, and clean reads were aligned to the reference genome using STAR (v 2.7.10b) [44] allowing no more than 3 mismatches. The raw read counts for each gene were extracted using featureCounts (v 2.0.3) [45], and the gene expression level was normalized by transcripts per million (TPM) using TPMCalculator [46]. To check the sample correlations, the principal component analysis (PCA) was conducted using the top 500 variable genes.

For identifying the differentially expressed genes (DEGs) between sexes, the raw read count matrix was imported in DEseq2 [47] R package. The pair-wise comparison was performed between female and male embryos, and DEGs were determined using the following criteria: |Fold Change (FC) >1.5| and FDR < 0.1. To remove the effect of potential sample batches, the sample collection time was included as a covariate in differential expression analysis. Additionally, an unsupervised hierarchical clustering was applied to the whole DEG list to highlight gene expression pattern between sexes, and the heatmap was generated using ComplexHeatmap [48] R package. The gene ontology (GO) and Kyoto Encyclopedia of Genes and Genomes (KEGG) pathway enrichments of sex-biased genes were then performed using clusterProfiler (v 4.12.6) [49] R package.

### Functional analysis of sex-biased regulators

To identify the sex-biased transcription factors (TF), we downloaded the bovine TF and cofactor list from AnimalTFDB (v 4.0) [35], and overlapped them with the DEGs. The expression pattern of RNA-binding proteins (RBPs) was assessed in bovine blastocysts through retrieving a comprehensive list of 1542 manually curated RBPs from human research [50] and converting them into bovine orthologs, and the sex-biased RBPs were obtained by overlapping with DEGs. We also checked the protein-protein interaction (PPI) networks for the DEGs using the Search Tool for the Retrieval of Interacting Genes (STRING, v12.0, https://string-db.org/), and proteins with interaction score < 0.4 were removed from our results. The PPI network was visualized with the Python hiveplotlib module to show the interaction patterns.

To check if the DEGs were significantly enriched in specific gene groups or genomic regions, we created six gene groups, including protein coding genes, TF, TF cofactor, and genes located on autosome, X and Y chromosomes. Fisher exact test was used to determine if the sex-biased genes were significantly more or less enriched in specific gene groups. In addition, we defined the promoter regions as ± 1 kb around the transcription start site (TSS) and performed motif searching using homer [51] to predict the upstream regulators of the DEGs. The motif results were summarized and clustered based motif sequence similarity using JunJunZai R package [52].

### Dosage compensation analysis

To evaluate the relationship between DEGs and dosage compensation in sex-biased early embryo development, the X chromosome to autosome (X:A) expression ratio was calculated for female and male embryos. The pairwiseCI R package was used to get a 95% confidence interval for the ratio of median of X to the median of A as in previous research [53]. 1,000 bootstrap replicates were included in the analysis where sampling from the original data was done with replacement and stratified by the group variables. Moreover, the X:A ratio was calculated for three gene categories, including expressed genes (TPM >1), DEGs, and non-DEGs between sexes. To remove the effects of non-unique mapped reads on gene expression, we further excluded the paralog genes in the bovine genome and calculated the X:A ratio for each of the defined gene categories. We also checked the distribution of DEGs on the X chromosome, and the gene density was calculated in a 1Mb window, which was visualized using RIdeogram [54] R package.

### Sex-biased alternative splicing event identification

To characterize the alternative splicing patterns in bovine early embryos, we followed the official protocol of rMATS-turbo [55] to obtain the sex-biased alternative events, including skipped exon (SE), alternative 5’ splice sites (A5SS), alternative 3’ splice sites (A3SS), mutually exclusive exons (MXE), and retained intron (RI). Briefly, rMATS-turbo utilized the widely used percentage spliced in (PSI) metric to quantify alternative splicing, which represents the percentage of transcripts that include a specific exon or splice site, as calculated from RNA-seq read counts supporting specific exons or splice junctions, normalized by the effective lengths of distinct transcript isoforms, then a generalized linear mixed model was applied to identify differential alternative splicing events between two groups. The statistically significant events were further selected with the following criteria: average RNA-seq read count >= 10 in both sample groups, filtering out events with average PSI value ranging from 0.05 to 0.95 in both sample groups, false discovery rate (FDR) <= 0.01, and between-group PSI value difference |ΔPSI| ≥ 0.05. The significant SE events between sexes were visualized using rmats2sashimiplot [56] and the potential RBP binding sites were predicted using rMAPS2 [57]. The functional enrichment of genes associated with differential alternative splicing (DSGs) was performed using clusterProfiler. Additionally, overlapping analysis was performed between DEGs and DSGs.

### Relationships between alternative splicing and known protein-coding domains

To explore if the differential alternative splicing (AS) events were associated with a switch between protein-coding isoforms and check if the SE, which was the most abundant AS event type in our analysis, can disrupt known protein domains, similar methods in a previous study were used [58]. Firstly, the coding sequence (CDS) was extracted from each isoform in the reference genome annotation and translated into corresponding amino acid sequence. Next, the known protein-coding domains were obtained from the PFAM database [59] and the amino sequences were used to query the PFAM database for selecting high-confidence amino acid alignment to determine protein-coding domains in each isoform. The high-confidence protein-coding domain alignments were defined with the following criteria: domain score > 10, domain E-value < 0.01, sequence alignment E-value < 1e-5, accuracy (hmmscan metrics) >= 0.8, and the proportion of the sequence domain aligned >= 0.9. To establish the relationships between SE events and protein-coding domains, the protein-coding domain locations were translated to genomic coordinates. Finally, the protein-coding domains were determined to be affected if they were overlapped with skipped exons, and enrichment analysis was conducted for the genes containing the affected protein-coding domains to check the potential functional changes.

### Long-read sequencing and differential isoform expression analysis

To capture the embryonic isoform diversity, pooled (n=25) 4-cell embryos were analyzed using PacBio long-read transcriptome sequencing. Briefly, 4-cell embryos were collected at 44 hours post insemination (hpi). RNA was extracted using PicoPure™ RNA isolation kit (KIT0204, Applied Biosytems™) with DNase treatment (RNase-Free DNase Set, Qiagen), ISO-Seq® express 2.0 kit for cDNA synthesis, SMRTbell® prep kit 3.0 for library preparation, and HiFi sequencing on SMRT Cell of a Revio platform (Genomics Resources Core Facility at Weill Cornell).

Long-read sequencing data were processed using the Iso-Seq pipeline [60]. Briefly, the high quality HiFi reads (predicted accuracy ≥ Q20) were processed with lima to conduct primer removal, and cDNA barcode identification. The poly(A) tail trimming and concatemer removal were conducted with isoseq refine. Then the isoform consensus clustering analysis was performed using isoseq cluster2, and the output isoforms were aligned to the bovine reference genome using pbmm2 [61]. Unique isoforms were obtained using isoseq collapse and 5’ degraded isoforms were removed using cDNA_Cupcake [62]. Finally, SQANTI3 (v 5.2.1) was used to classify isoforms with default parameters.

Next, we added the novel isoforms identified by long-read to the bovine reference genome. Using this new annotation, isoform expression was quantified using Salmon [63], and significantly differential expressed isoforms were identified using the swish method of fishpond [64] with q-value < 0.05. Additionally, the sex-biased exon usages (padj < 0.05 and log2FC >2) were examined using DEXSeq [65] with default settings.

### Validation of differential expression genes

Real-time quantitative polymerase chain reaction (RT-qPCR) was used to validate the result of DEG identified by RNA-seq. 10 sex-biased DEGs were selected based on their fold change, p-value, and chromosome location. Five were upregulated in females (3 on the X chromosome and 2 on autosomes) and five were upregulated in males (3 on the Y chromosome and 2 on autosomes). Primer sequences were listed in **Table S2.**

A 10ul qPCR reaction contained 2 ng of cDNA, iTaq^TM^ Universal SYBR^®^ Green Supermix (Bio-Rad), and 0.5 uM of primers. The qPCR conditions were: 95°C for 3min; 30 cycles of 30 s at 95°C, 58 °C for 30s, and 72 °C for 45s; and 72 °C for 10 min. Relative expression was calculated using the 2^-ΔΔCT^ method with *GAPDH* and *H2AZF* used as reference genes. For evaluating the relationships between qPCR and RNA-seq results, we fit a linear regression model based on the relative expression fold changes derived from these two methods for the selected genes.

### Validation of alternative splicing events

Seven genes that with differentially exon skipping events between sexes were validated by PCR. A 10 uL of reaction contains 4 ng of cDNA, 5 uL of One*Taq*® Quick-Load® 2× Master Mix (M0486L, New England Biolabs), and 5 pmol each of forward and reverse primers. PCR was conducted with the condition as follows: 15 minutes at 95°C; followed by 35 cycles of 30 seconds at 95°C, 30 seconds at 58°C, and 45 seconds at 72°C; lastly 72 °C for 10 min. PCR products were visualized by gel electrophoresis using a 2% agarose-TAE gel. Primers were designed to span the predicted skipping exon and were listed in **Table S3**.

## Results

### Sex-specific development paces

On Day-7 (168 hpi), expanded and hatched blastocysts were collected, while non-expanded blastocysts were kept in the culture medium for another 24 h. On Day 8 (192 hpi), non-expanded, expanded, and hatched blastocysts were all collected. Expanded blastocysts were further divided into two even halves for sex determination or transcriptomic sequencing. Each half contained both the inner cell mass and trophectoderm (**Figure 1A**). In total, 444 blastocysts were collected. 412 of them were successfully sexed by the genomic PCR with sex-related primers and visualized on electrophoresis gel imaging (**Figure 1B**). A significant male-biased development was observed among the collected blastocysts, except for Day 7 hatched embryos (**Table S4**). On Day 7, 107 out of 155 expanded blastocysts were males (sex ratio = 2.23, p < 2.15e-06), whereas no significant sex ratio difference in hatched blastocysts due to few embryos hatched on Day 7 (8 males vs. 4 females, sex ratio = 2, p = 0.25). On Day 8, all developmental stages exhibited a significant male-biased development, including non-expanded (80/99, sex ratio = 4.21, p < 8.75e-10), expanded (91/112, sex ratio = 4.33, p < 3.73e-11), and hatched blastocysts (29/34, sex ratio = 5.80, p < 3.86e-05). These results aligned with the previous studies reporting a higher proportion of male expanded and hatched *in vitro* produced blastocysts on Days 7 and 8 in cattle [1, 13, 66, 67]. Moreover, male-specific genes on the Y-chromosome showed significant low/no expression in female samples, confirming a high accuracy of sex identification (**Figure 1C**).

### Summary statistics of the RNA-seq data

5 male RNA-seq samples (5 pools of 10 half-cut male expanded blastocysts) and 6 female RNA-seq samples (6 pools of 12 half-cut female expanded blastocysts) with known sex information determined by PCR were used for RNA-seq. RNA-seq was performed using the Illumina PE150 platform. Data quality and mapping information were summarized in **Table S4**. A total of 399,685,192 raw reads were generated in the 11 samples, averaging 36,335,017 reads per sample with an error rate of 0.04%. GC content of the samples ranged from 45.23% to 47.60%, while sequencing quality metrics Q20 and Q30 were above 89.94% and 83.16%, respectively. Unique read mapping rates to the bovine reference genome averaged 80.94%, indicating robust data quality for bioinformatic analysis.

### Identification of sex-specific differentially expressed genes (DEGs)

To assess transcriptomic variation and clustering between sexes, principal component analysis (PCA) was conducted. Using the top 500 most variable genes, samples grouped into two distinct clusters, consistent with our experimental design, confirming that female and male embryos have different transcriptomic profiles at the early embryo stage (**Figure S1**). Sex-specific transcriptomic profiles were expected due to the different sex chromosome compositions between males and females. Indeed, 837 DEGs were identified between the two sexes, with 231 upregulated DEGs and 606 downregulated DEGs in male expanded blastocysts when compared to female expanded blastocysts (**Figure 2A, Table S5**). Additionally, protein-protein interaction analysis revealed intensive interactions both within the DEG groups and across the non-DEGs (**Figure S2)**, which may help to establish the sex-specific regulatory networks during early embryo development (53). The two most differentially expressed genes *XIST* and *USP9Y*, located at the X chromosome and Y chromosome, were significantly upregulated in female and male expanded blastocysts, respectively. Among all the DEGs, 574 were located at the autosomes, 250 were located at the X chromosome, and 13 were located at the Y chromosome (**Table S6**).

**Figure 2.**
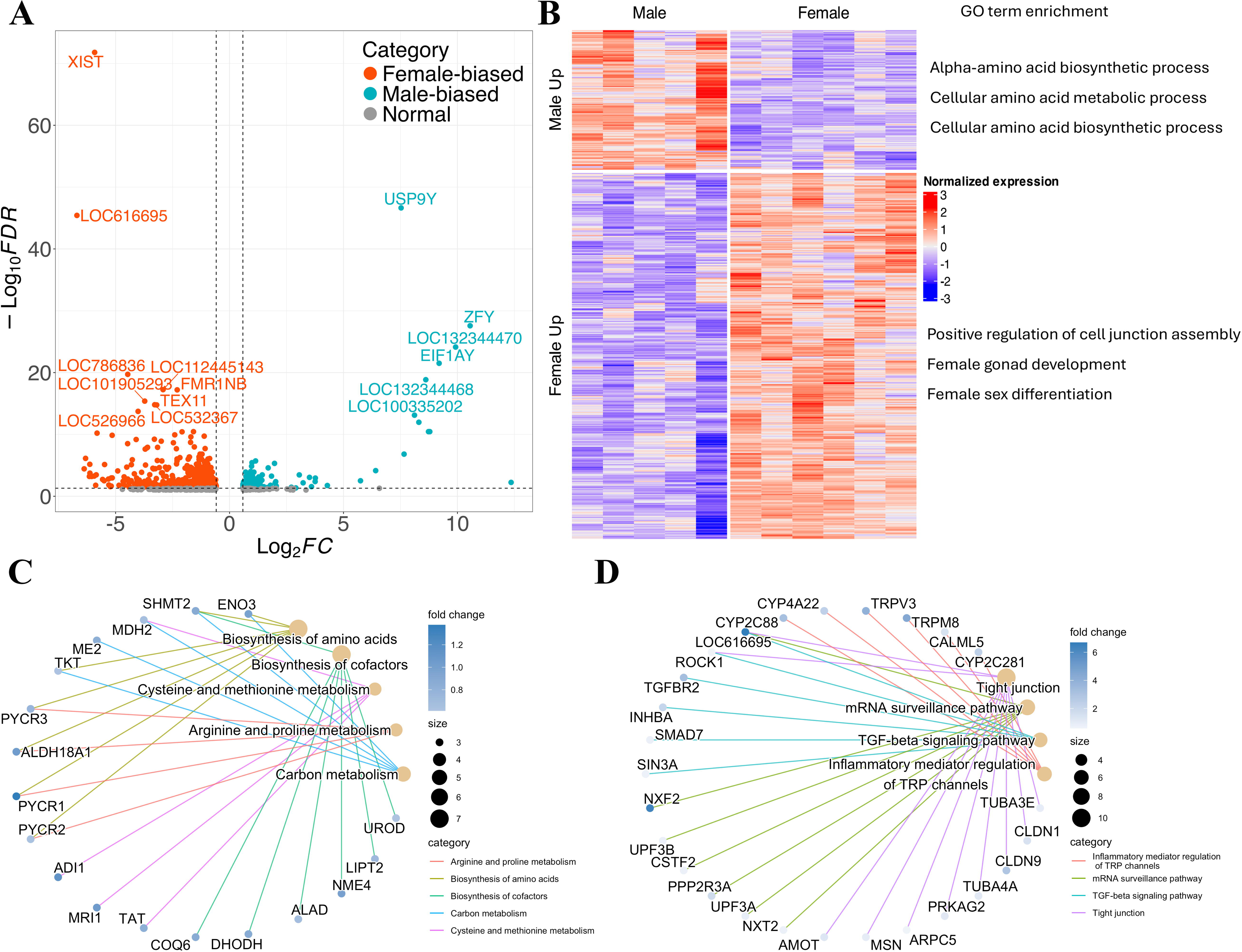
Sex-biased genes of bovine expanded blastocysts and functional enrichment. A) Volcano plot of all DEGs between male and female expanded blastocyst (BL). B) Heatmap and GO functional terms of sex-biased genes. C) KEGG pathways enriched in male-biased DEGs in expanded BL. D) KEGG pathways enriched in female-biased DEGs in expanded BL.

### Function and regulation of sex-biased DEGs

To further explore these sex-specific transcriptomic differences, we performed an unsupervised hierarchical clustering of the entire sex-biased DEG list (**Table S5**). The DEGs were grouped into two clusters, corresponding to female- or male-biased genes (**Figure 2B**). Gene ontology (GO) analysis showed that male-biased DEGs were primarily associated with metabolic processes, with top functions such as alpha-amino acid biosynthetic, cellular amino acid metabolic, and cellular amino acid biosynthetic process. In contrast, female-biased DEGs were enriched in processes such as female gamete generation, ovulation cycle process, positive regulation of cell junction assembly, and female gonad development (**Figure 2B**). Moreover, the enrichment results of KEGG analysis showed that male-biased DEGs were enriched in metabolism and biosynthesis pathways, while female-biased DEGs were associated with inflammatory responses, cellular activity, TGF-beta signaling, and tight junction (**Figure 2C and 2D**).

Examination of bovine known transcription regulators in sex-biased DEGs identified 68 sex-biased expressed TFs and cofactors, with 19 showing higher expression in males and 49 in female embryos, respectively (**Figure 3A**). Specifically, we observed a few male-biased pluripotency regulators, including *HMGA2, SOX21,* and *PRDM14*. Moreover, we converted the human RNA binding proteins (RBP) list to cow orthologs and identified 31 and 21 RBPs exhibiting female- and male-biased expression, respectively (**Figure 3B).** Next, we determined the genomic distribution of sex-biased genes, we found that female-biased genes were significantly enriched in protein-coding genes, TF cofactors, and X chromosomes, while male-biased genes were significantly enriched in protein-coding genes and Y chromosome (**Figure 3C**), indicating the distinct regulation of sex chromosomes in sex-specific early development

Next, we performed *de novo* TF identification by motif calling based on promoter regions of the sex-biased DEGs. This analysis revealed several sex-biased motifs with predicted upstream binding TFs (F**igure S3**). For example, the female-biased motifs were dominated by TFs such as *ETV, ELK, ELF, SP*, and *KLF* families, whereas the male-biased motifs included *KLF, SP, ETS,* and *ELF* TF families. Shared TF families between the two sexes included *PABPC1, NFY, ELK4, ELFs, SPs,* and *KLFs,* with *PABPC1* showing the most significant enrichment (**Figure S3**).

**Figure 3.**
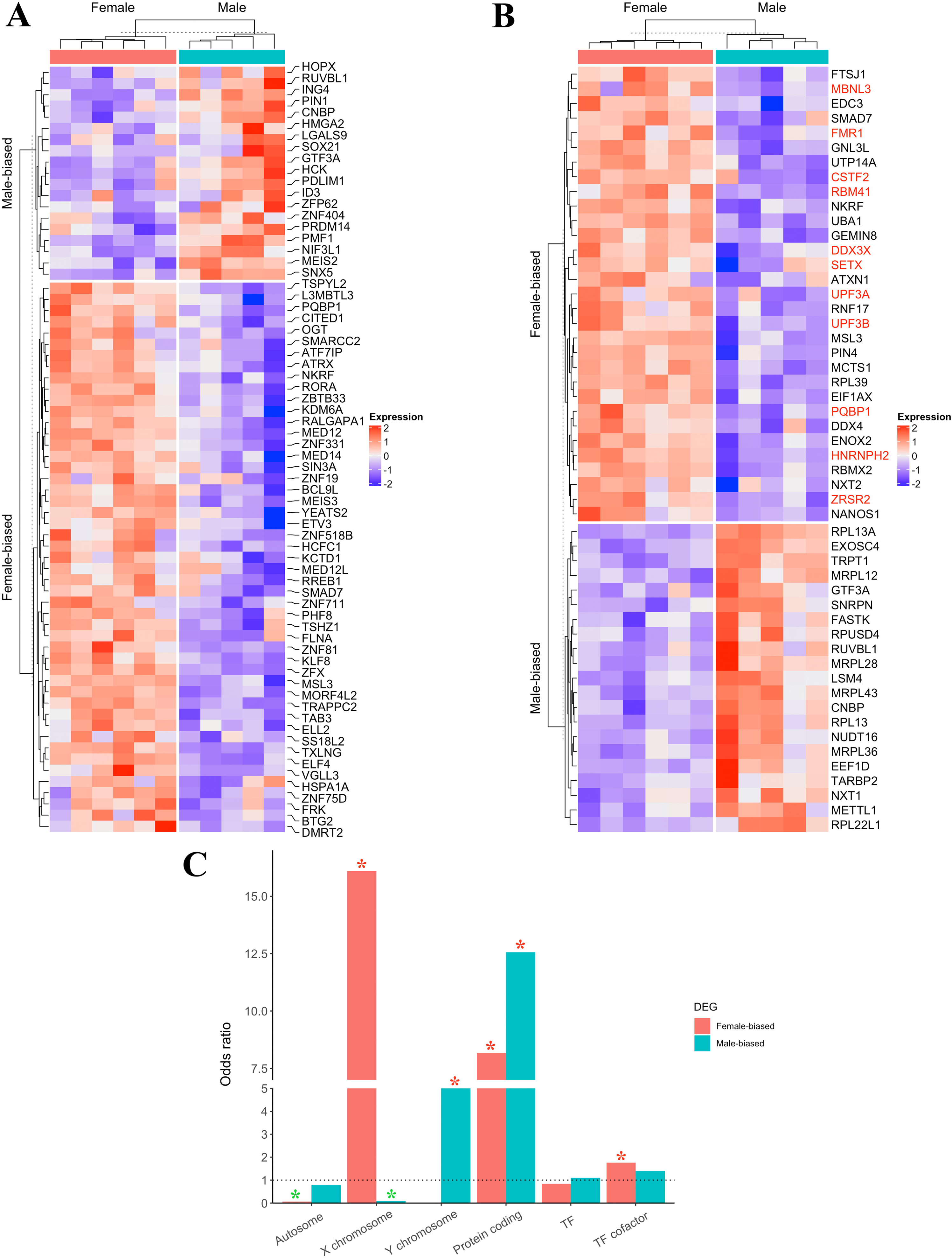
Heatmaps of Sex-biased transcription regulators and genomic distribution of DEGs. A) Sex-biased transcription factors and cofactors. B) Sex-biased RNA binding proteins (RNA splicing factors highlighted in red). C) Enrichment analysis of DEGs in different gene groups. Green asterisks mean significantly less enrichment (p-value < 0.01) and red asterisks means significantly over enrichment (p-value < 0.01).

### X-chromosome dosage compensation in sex-specific embryos

To investigate the X-chromosome dosage compensation at the blastocyst stage, X:A ratios were calculated in three categories: expressed genes (TPM>1), DEGs, and non-DEGs for both sexes (**Figure 4**). An X:A ratio of 1.0 or higher indicates complete compensatory upregulation of X-linked genes compared to autosomes, 0.5 suggests no compensation, and values between 0.5 and 1 reflect partial compensation [68]. In male embryos, the X:A ratios were consistently around 1 in expressed genes and slightly lower than 1 in male-biased DEGs (**Figure 4**), indicating a complete and incomplete dosage compensation, respectively. In contrast, the X:A ratios varied in female embryos, with the highest value (> 1.5) observed in female-biased DEGs, around 1.3 for the expressed genes, and close to 1 for non-DEG. These findings suggested that the dosage compensation levels differed among different subgroups of the X-linked genes, with female-biased X-linked DEGs being the primary drive of dosage.

**Figure 4.**
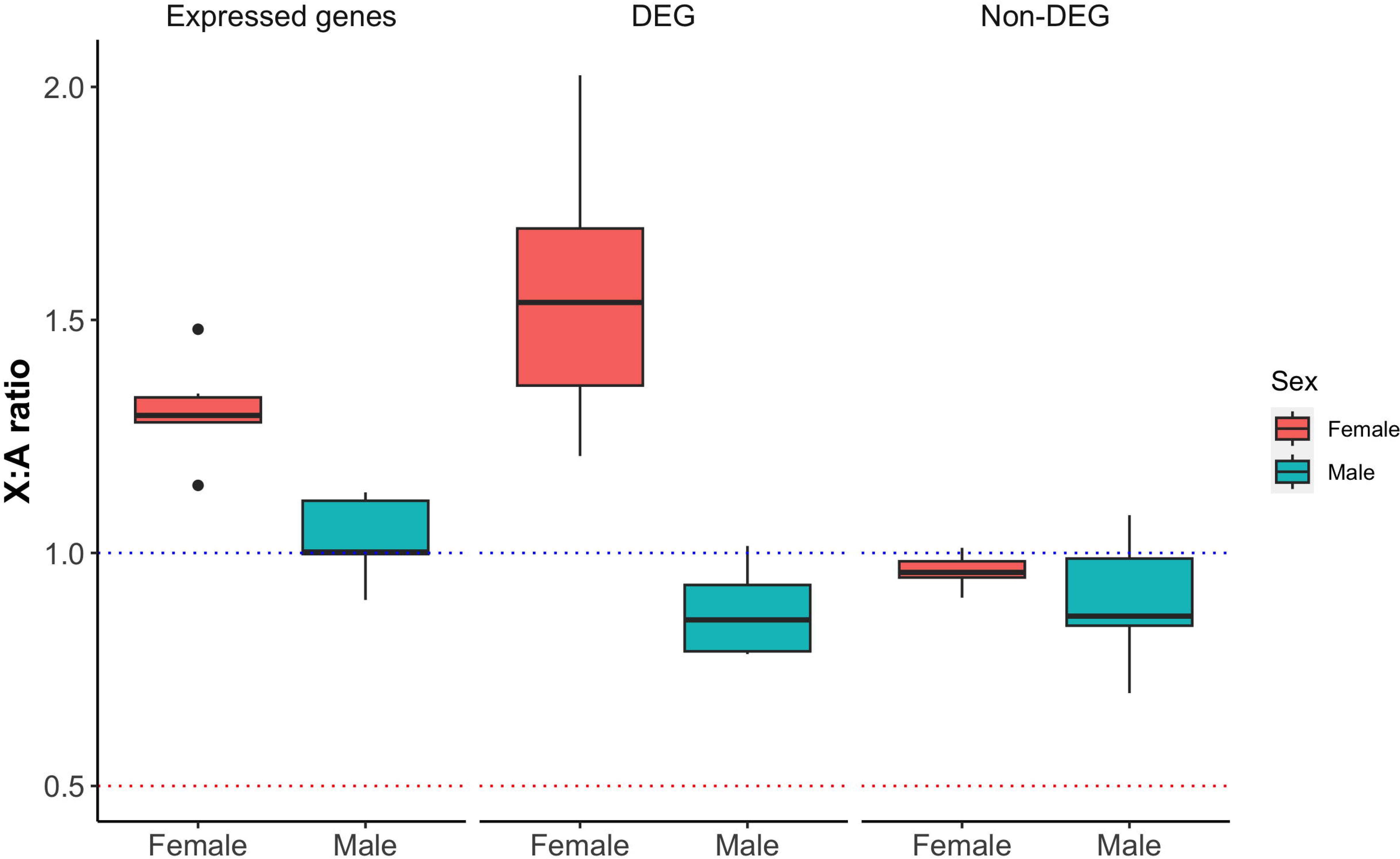
Gene dosage compensation. The X:A expression ratio was calculated using expressed genes (TPM >1), DEGs and non-DEGs in female and male expanded blastocysts.

Given the fact that the X chromosome is abundant with paralog genes in mammals [69, 70], which might affect the quantification of the X-linked genes, we recalculated the X:A ratio after excluding the paralog genes in the bovine genome. Interestingly, the X:A ratio was reduced when excluding the paralog genes, especially in the sex-biased DEGs group, further supporting the enrichment of paralog genes on the X chromosome contribute to the dosage compensation (**Figure S4**).

To further assess if the elevated expression of X-linked DEGs in females was associated with X chromosome inactivation (XCI) and located to specific XCI regulatory regions such as the X inactivation center, gene-rich areas, or pseudoautosomal regions (PAR) that escape XCI [69, 71], we compared the genomic distribution of the female-biased X-linked DEGs with all X-linked genes (**Figure S5**). The result showed a strong overlap of locations between female-biased genes and certain gene-rich regions along the X chromosome, however, these female-biased X-linked DEGs did not overlap with known PAR regions, which were located by the very end of bovine X [72]. Overall, our analysis suggests the upregulated expression of female-biased DEGs was not linked to specific XCI regulatory regions of the X chromosome.

### Alternative splicing in sex-specific embryos

Next, we evaluated the contribution of alternative splicing to sex-specific early embryonic development. Using rMATs, we identified 27,875 AS events, with 1,555 showing significantly sex-biased difference (FDR<=0.01; delta PSI value ≥0.05) across five different event types: skipped exon (SE), alternative 5’ splice sites (A5SS), alternative 3’ splice sites (A3SS), mutually exclusive exons (MXE), and retained intron (RI) (**Figure 5A, Table S7**). Among the five types, SE event was the most dominant AS category between sexes (**Figure 5B**), with 281 SE events being female-biased and 475 SE events being male-biased. For example, *ZMYND11*, a gene involved in epigenetic regulation [73], exhibited significant high exon inclusion in male embryos (**Figure 5C**), while *CEP78* showed a significantly high exon inclusion level in female embryos (**Figure 5D**). Functional enrichment analysis of sex-biased AS events associated genes revealed that the female-biased AS genes were enriched in amide and peptide biosynthetic processes, while the male-biased AS genes were enriched in mRNA processing and metabolic processes (**Figure 5E**). Moreover, overlapping analysis between sex-biased AS genes and sex-biased DEGs identified 59 AS genes that also exhibited significant differential expression at the gene level (**Figure 5F**). GO analysis of these overlapping genes revealed their association with translation regulation, protein metabolic process, mitochondrial respiratory chain, and oxidoreductase activity, all of which played essential roles in early embryogenesis, indicating their potential roles in sex-specific developmental regulation (**Table S6**).

**Figure 5.**
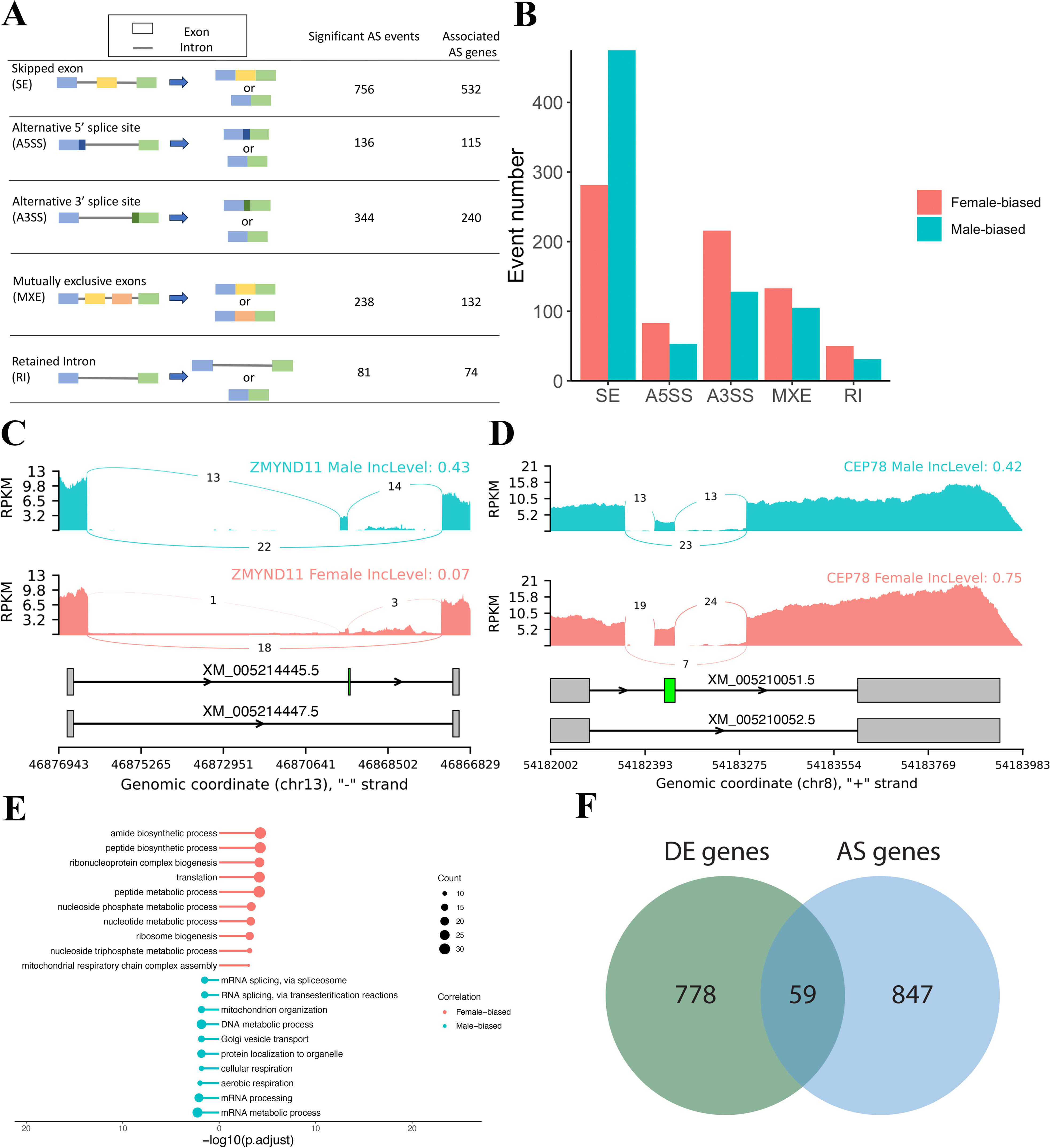
Sex-biased alternative splicing events. A) Five categories of alternative splicing (AS) events examined by rMATs, with numbers of Sex-biased differential AS events (right panel, FDR <=0.01 & delta PSI value (≥0.05) and AS associated gene. B) Event number of female- and male-biased differential AS events. C) Illustration of a male-biased SE event in Z*MYDN11*, with a higher inclusion level of the transcripts that contain the skipped exon in male blastocyst sample. D) Illustration of a female-biased SE event in *CEP78*, with a higher inclusion level of the transcripts that contain the skipped exon in female blastocyst sample. E) GO term analysis of all sex-biased AS associated genes. F) Venn diagram of overlapped genes between differentially expressed gene list and sex-biased AS event associated gene list.

Furthermore, we investigated the relationships between the sex-biased SE events and two known splicing factors, *FMR1* and *HNRNPH2,* both of which were also DEGs. In female-biased SE events, *FMR1* showed significantly increased binding potential downstream of the alternative exons, whereas in male-biased SE events, *HNRNPH2* exhibited significantly increased binding potential upstream of the alternative exons, suggesting distinct sex-specific AS regulatory mechanisms (**Figure S6**).

Finally, to determine whether alternative exon usage overlaps with protein-coding domains and contributes to protein diversity, we mapped the known protein domains to sex-biased SE events. A total of 280 and 438 skipped protein domains were identified in female and male embryos, respectively (**Table S8**). Functional enrichment analysis of associated genes exhibited skipped protein domains revealed that energy metabolism and protein process were enriched in female embryos, while the transport and catabolism process were enriched in male embryos (**Table S8**).

### Differential isoform level expression in sex-specific embryos

Using long reads data, we identified 286,711 unique isoforms that mapped to 14,084 novel genes and 14,353 annotated genes. Through comparison with the cow reference genome, the long-read isoforms were classified into different categories, including full splice match (FSM), incomplete splice match (ISM), novel in catalog (NIC), novel not in catalog (NNC), intergenic, antisense, fusion and genic genomic (**Figure S7A**). In total, we obtained more than 140,000 novel isoforms (NIC and NNC categories) from the early embryos (**Figure S7B**). To assess the isoform features, we found more than 50% of the genes only have one isoform, around 10% of the genes have 2-3 isoforms, and the rest of the genes have at least four isoforms (**Figure S7C**). Additionally, more than 90% of the isoforms contained the canonical splicing sites and had relatively high short-read coverage associated with the splice junctions, suggesting good quality of the long-read data (**Figure S7D**).

To capture a comprehensive view of the embryonic isoforms, we added the novel isoforms to the cow reference genome annotation and performed differential isoform analysis. A total of 1,151 differentially expressed isoforms were obtained in this study, including 514 female-biased isoforms and 637 male-biased isoforms, which were associated with 1,017 genes (**Table S9**). According to the functional enrichment of the differential isoform-associated genes, we showed that negative regulation of DNA-templated transcription and RNA biosynthesis were enriched in female embryos (**Figure 6A**), and the peptide metabolic processes were enriched in male embryos (**Figure 6B**), aligning with the slow development pace observed in female embryos. Overlapping analysis identified 419 genes that had differentially expressed isoforms and were also sex-biased DEGs, as well as 478 genes that only contained differentially expressed isoforms (**Figure 6C, Table S10),** including *APOE, KDM6A,* and *GOLM2* (**Figure S8**). Additionally, 160 genes overlapped between sex-biased AS events and sex-biased isoforms, while only 19 overlap genes between sex-biased AS events and sex-biased DEGs, suggesting that sex-biased AS events contribute more to differential isoform expression than to sex-biased DEGs (**Figure 6C**). Furthermore, analysis of differential exon usage between sexes revealed 7 exons exhibit biased usage in female or male embryos (padj <0.05 and log2FC >2). These exons were associated with 5 genes, including *DENND1B*, *DIS3L2*, *DOCK11*, *IL1RAPL2*, and *ZRSR2Y* (**Figure S9 and Table S11**), which may have sex-specific regulation in early embryo development.

**Figure 6.**
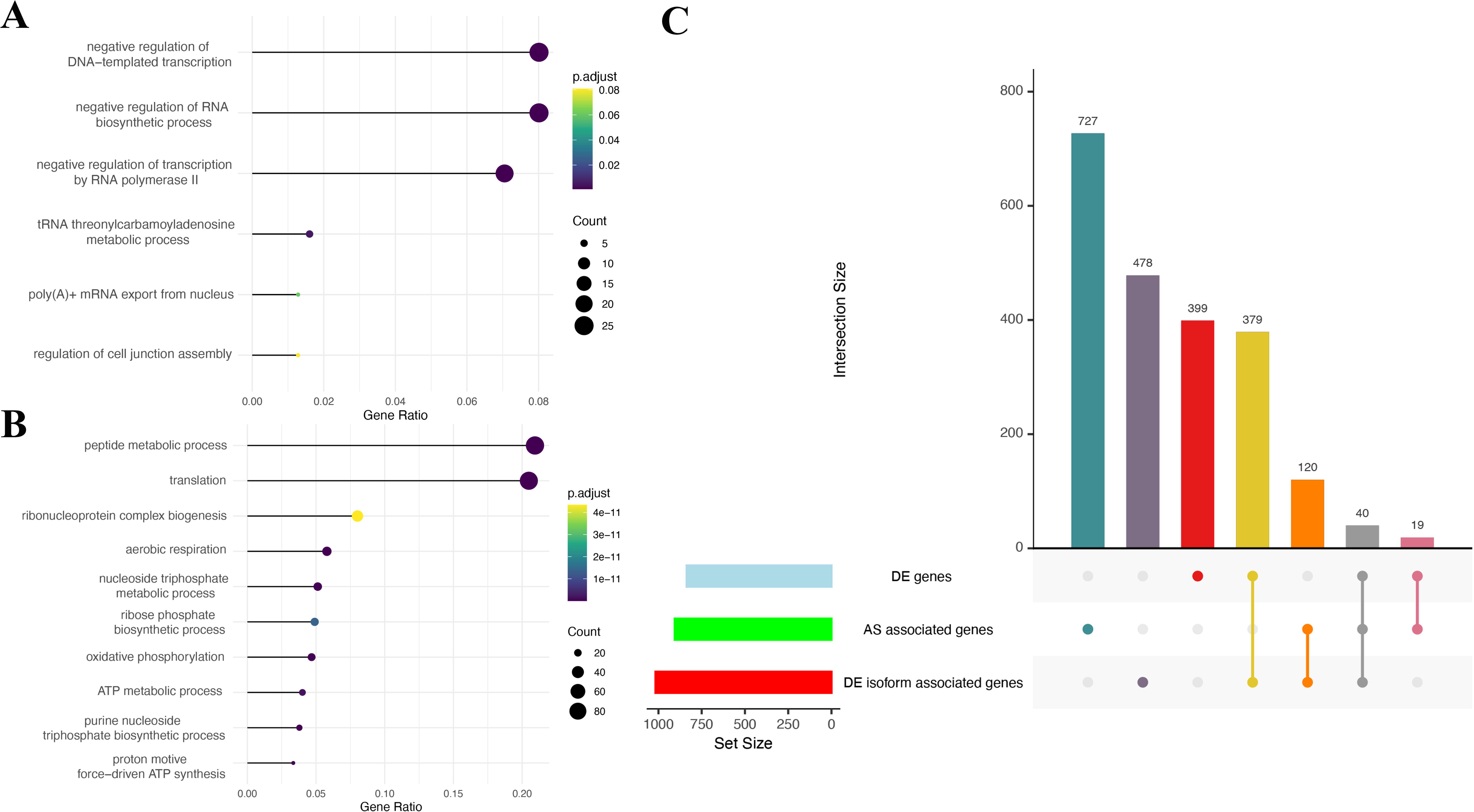
Functional enrichment of sex-biased isoforms and associated genes. A) GO functional enrichment of female-biased isoforms. B) GO functional enrichment of male-biased isoforms. C) Overlap genes among DEGs, sex-biased AS associated genes and sex-biased isoform associated genes.

### Validation of sex-biased DEGs and alternative splicing events

For validating the gene expression pattern between sexes from RNA-seq, we selected the top sex-biased genes (log2FC > 1.5) and quantified their expression using PCR methods, including female-biased autosomal genes (*GGNBP1*, *SVIP*), male-biased autosomal genes (*MIOX*, *NNAT*), female-biased X-linked genes (*MAGEB1*, *MAGEB16*, and *XIST*), and male-biased Y-linked genes (*DDX3Y*, *EIF1AY*, and *ZRSR2Y*). The relative expression fold change among all examined genes exhibited similar trends between RNA-seq and PCR results (R^2^ =0.83, **Figure 7A**), indicating the robustness of our RNA-seq analysis.

**Figure 7.**
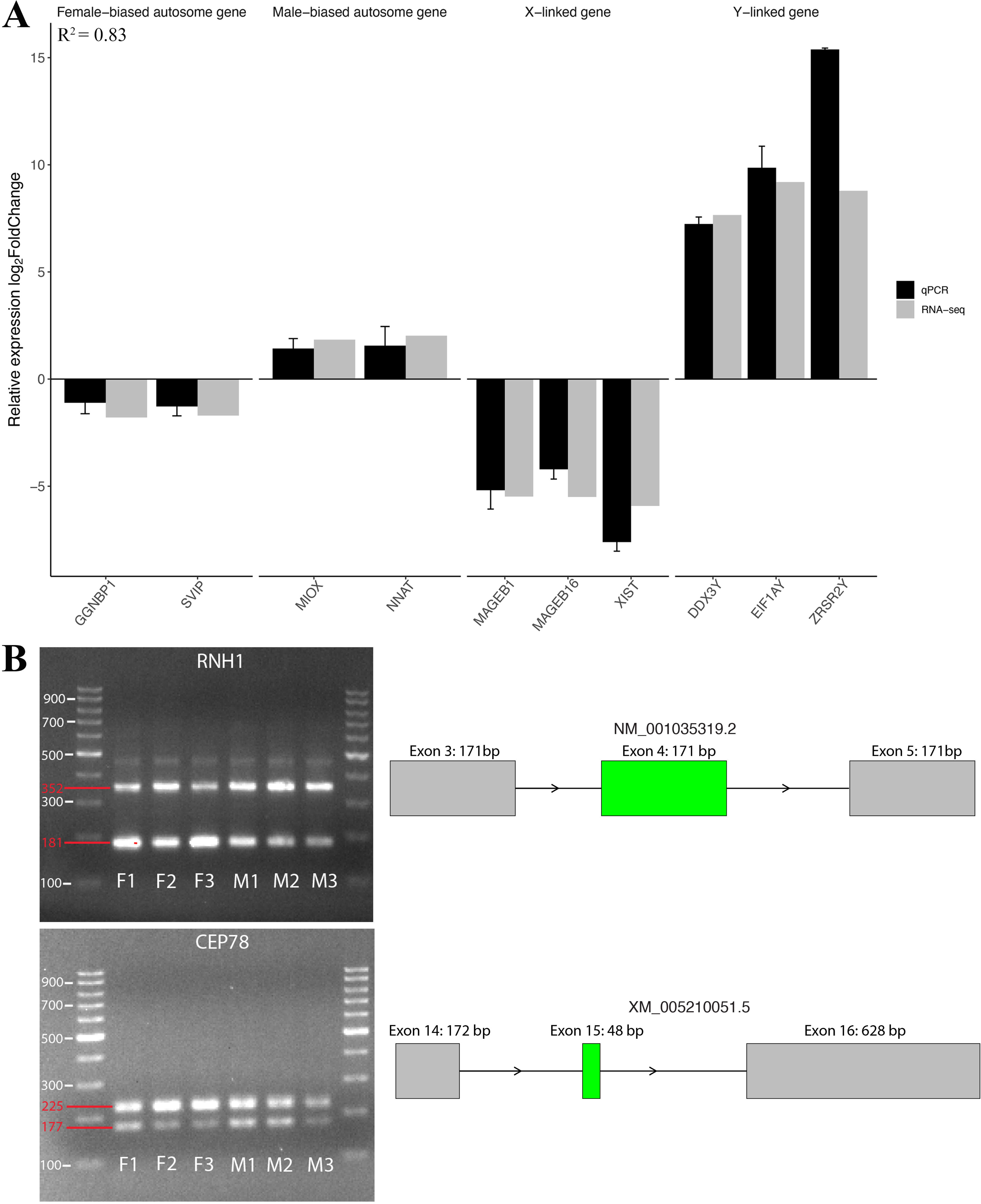
Validation of DEGs and AS events. A) Validation of DEGs with qPCR. B) Agarose gel of two selected alternative splicing associated genes (RNH1 and CEP78). Primers were designed to span the predicted skipping exon. The upper band contains the transcripts that include the skipped exon, while the lower band contains the transcripts that do not include the skipped exon. F1, F2, and F3 are female embryos; while M1, M2, and M3 are male embryos.

To validate the sex-biased AS events, we selected five significantly differential SE events and designed specific primers to amplify the target regions of the alternative exons, including one female-biased SE in *CEP78* and four male-biased SEs in *RNH1*, *CEP350*, *ZMYND11*, and *WWC2*, respectively **(Figures 7B and S10)**. Gel electrophoresis results showed that in male embryos, the exon inclusion band (352bp) of *RNH1* was brighter than the exon-skipped band (181bp), while in female embryos, the exon inclusion band (225bp) of *CEP78* was more abundant than the exon-skipped band (177bp), consistent with the sex-biased exon inclusion level observed in RNA-seq data (**Figure 7B**). Further confirmation showed that the observed PCR product length differences were due to the skipped exons, and the exon inclusion level calculated from PCR gel electrophoresis aligned with the RNA-seq calculated results (**Figure S11**).

## Discussion

Sex differences are a widely studied topic in developmental, nutritional, and medical research, as the distinct composition of sex chromosomes influences hormone synthesis, brain function, immune regulation, and body structures, causing males and females to respond differently to the same stimuli. While many of the differences become apparent only after the formation of sex glands and the onset of hormone production, embryos already exhibit sex-specific differences during the preimplantation period, even before sex-specific hormonal regulation [21].

In this study, we investigated the transcriptomic differences between male and female bovine embryos in early development. Using short-reads RNA-seq and long-reads ISO-seq, we identified sex-biased DEGs, AS events, and isoform-level expression patterns. Male embryos exhibited faster development, driven by upregulated metabolic pathways, while female embryos showed slower development associated with negative regulation of transcription and RNA biosynthesis. Additionally, distinct X-chromosome dosage compensation was observed, with an X:A ratio greater than 1 in female embryos, suggesting both X-chromosome upregulation (XCU) and incomplete X-chromosome inactivation (XCI). Overall, these findings provide new insights into the regulatory mechanisms underlying sex-specific embryogenesis.

### Sex-specific embryonic development in different species

Sex differences in early embryos, observed before gonadal formation and hormone synthesis, are well-documented across species. Previous studies in cattle have shown that *in vivo* developed male blastocysts retrieved from the cows exhibit faster development, higher proliferating cell numbers, and elevated expression of pluripotent genes [31]. Similarly, our study demonstrated that the *in vitro* produced bovine male blastocysts developed faster than female embryos, consistent with previous findings in IVF embryos in cattle [1, 13, 14, 17]. This phenomenon is conserved among other mammals, including humans [3–9], mice [10–12], pigs [15, 16], and sheep [17], as well as in porcine *in vivo* embryos [15]. However, the observations in humans are more complex. While some studies suggest no difference in live birth sex ratio following IVF embryo transfer [74], others have observed accelerated male development in ICSI-produced embryos but not in IVF embryos [22]. These discrepancies may arise due to variations of embryo origins, culture conditions, and study designs, as human studies often involve embryos from oocyte or sperm donors with fertility issues.

Interestingly, environmental factors such as the composition of *in vitro* culture medium and culture conditions also significantly influence sex-specific development. For example, glucose supplementation in the culture medium was found to accelerate male development and increase cell numbers in male bovine blastocysts [21, 22, 31]. Moreover, cultures supplemented with serum or plates coated with granulosa cell monolayer enhance the male-to-female ratio at Day 6 and Day 7, indicating faster cleavage in male embryos under these conditions [1]. In contrast, female embryos exhibit greater resilience and higher survival rate compared to males under heat stress, both *in vivo* and *in vitro* culture [75]. These studies highlighted the sex-specific genome regulation in driving the differences in development speed and response to different culture conditions. However, the underlying regulatory mechanisms remain incompletely understood.

### X dosage compensation in sex-biased embryogenesis

Transcriptional sexual dimorphism in early embryos is influenced by factors such as the expression of X/Y-linked genes, XCI escapees, and interactions between sex chromosomes and autosomal genes [21]. In bovine somatic tissues, studies have reported a complete X dosage compensation for “dosage-sensitive” (ubiquitously expressed) genes with X:A ratios around 1 and no sex difference [69]. In bovine *in vivo* embryos, X:A ratios vary dynamically: they start at 1 in maturated oocytes, decrease in zygotes with an inactive paternal X, rises to peak levels (X:A >1) during ZGA and reduce to around 1 after the morula and blastocyst stages, where it stabilizes [69]. Interestingly, in IVF embryos, the X:A ratio showed distinct patterns compared to *in vivo* embryos with X:A ratio exceeding 1 in IVF blastocysts [69]. These findings reflect the intricate regulation of X dosage during bovine early embryo development and the influences of the embryonic development environment.

According to Ohno’s hypothesis of X dosage compensation [76, 77], in somatic cells, the XCU compensates for the silencing effects of XCI by upregulating genes on active X in females, maintaining an X:A ratio close to 1. Consistent with this, our study found that the X:A ratio for expressed genes was around 1.3, while for female-biased DEGs, it was 1.5 in female blastocysts. This upregulation of X-linked genes in female embryos is also consistent in human embryos [32, 33]. The elevated X:A ratio of the expressed and female-biased DEGs likely reflects the XCU coupled with an incomplete XCI, as *Xist* expression is initiated at the morula stage and may not yet be completely established in blastocysts. Specifically, the upregulation of female-biased DEGs may result from XCU on the active X chromosome (Xa) while some genes remain active on the Xi due to incomplete XCI, or they are classified as XCI escapees, contributing to the elevated X:A ratio.

In mice, XCU has also been observed during early embryogenesis, and males achieve XCU upon ZGA while there are two waves in females: the first in response to imprinted XCI after ZGA and until blastocyst stage, then the second during random XCI at post-implantation stage [78, 79], both mediated by *Xist* [80, 81]. Impaired XCI leads to a subsequently skewed sex ratio of embryos with female-biased peri-implantation defects [82]. In mouse IVF embryos, this skewed sex ratio has been linked to reduced expression of XCI regulators, such as *Rnf12* or *Xist*, which can be rescued by *Rnf12* overexpression [82]. These findings suggest that the incomplete XCI in female blastocysts may partially contribute to a slower development rate and the skewed sex ratio in early development. However, further studies are needed to validate this hypothesis.

### Sex-biased DEGs in early embryos

Cellular programming, activity, epigenetic modifications, and gene transcription are closely linked to metabolism, with metabolites signaling primarily influencing epigenetics and gene transcription, while metabolism pathways are largely regulated by post-translational mechanisms [83]. During oocyte maturation and early embryo development, maturation and initial cleavages rely primarily on maternal deposits, such as mitochondrial, mRNAs, proteins, and lipids, whereas later divisions depend more on embryonic synthesis and nutrients from the developmental environment [84–86]. Upon blastocyst formation, more oxygen, nutrients, and energy consumption are required for cell growth and division [84]. In drosophila embryogenesis, abnormal high dNTP levels induced by expression of feedback-insensitive ribonucleotide reductase accelerate nuclear cleavages while reducing zygotic transcription, resulting the failure in early gastrulation [86]. Additionally, nutrient depletion, such as glucose or phosphate removal from culture media, often leads to hamster or human embryo arrest *in vitro* [87–89]. The inadequacies of the *in vitro* culture condition compared to the *in vivo* environment contribute to delayed embryo development in hamsters [90]. These findings suggest that a more active metabolism correlates with faster early embryonic development.

In our study, male-biased DEGs were enriched in amino acid biosynthetic and metabolic processes (e.g. *ENO3*, *PYCRs*), mitochondrial structures (e.g. *COQ6*, *MRPL43*, *SHMT2*), and carbon metabolism (e.g. *SHMT2*, *ME2*, *TK*), suggesting a higher metabolic activity in male blastocyst. This finding is consistent with a microarray study, which reported high expression of genes related to metabolism, mitochondrial regulation, and cell cycle processes in male blastocysts compared to females [29]. Among amino acids crucial for cell growth and embryo development, proline enhances mitochondrial function and cell proliferation [91, 92], arginine promotes trophoblast development [93], and methionine, essential for DNA methylation, is linked to pluripotency maintenance [94]. Additionally, increased mitochondrial activity is associated with aster transcription, translation, and proliferation in human cells culture [95]. Therefore, enriched amino acid and carbon metabolism may accelerate blastomere cleavage in males, contributing to earlier development to the blastocyst stage.

In contrast, female-biased DEGs were associated with female-specific developmental processes, including gamete generation, gonad development, and ovulation cycles, with involvement in pathways such as mRNA surveillance, TGF-beta signaling, and inflammatory mediation. While gonad development is not initiated at this stage, the upregulation of female-specific development genes suggests a sex-specific developmental difference. Moreover, the mRNA surveillance pathway plays a critical role in detecting and degrading abnormal mRNA, and the disruptions in this pathway have been linked to lower embryo viability and infertility in mice [96]. Additionally, TGF-beta signaling, which regulates pluripotency in embryonic stem cells [97], is highly expressed at the 4- to 8-cell stages but declined at the blastocyst stages in cattle [98, 99]. Dysregulation of this pathway way leads to elevated TGF-beta signaling levels and results in bovine blastocysts degeneration [100, 101]. These findings suggested that the *in vitro* produced bovine female blastocysts exhibit female-specific regulatory mechanisms during early development, whereas male blastocysts show higher metabolic activity. However, a transcriptomic study has reported a higher metabolic activity in *in vivo* produced bovine female blastocysts [30], indicating that the developmental environment may contribute to this difference.

According to previous studies, faster cleavage can be associated with enhanced metabolic activity, but it does not always directly indicate better embryo quality. For example, in mice, faster-developed IVF embryos exhibit higher rates of imprinting errors, such as imprinted methylation loss of *H19* and *Snrpn* [102], whereas slower-developed embryos are more prone to development arrest. Both aberrantly developed embryos show higher aneuploidy rates compared to moderate cleavage rate IVF embryos, which more closely resemble the imprinted pattern of *in vivo* embryos, leading to lower implantation and pregnancy rates [103]. These findings indicate that faster-developing blastocysts are not necessarily of higher embryonic quality.

Beyond gene functional analysis, upstream transcriptional regulators of DEGs also exhibit sex-specific differences. Among the male-biased transcription factors, *HMGA2, SOX21*, and *PRDM14*, which are key regulators of pluripotency maintenance [104–106], align with enriched pluripotency pathways in males. Among the sex-biased RBPs, 11 RBPs were known to regulate alternative splicing (AS) in humans [107–113], including *MBNL3, FMR1, CSTF2, RBM41, DDX3X, SETX, UPF3A, UPF3B, PQBP1, HNRNPH2, and ZRSR2*. Notably, *FMR1* and *HNRNPH2* are two known splicing factors associated with fragile X syndrome and X-linked development disorders [114, 115], they exhibited significant female-biased expression. Moreover, the female-biased RBPs showed enrichment for AS regulation, supporting the observation that AS profiles are distinct between female and male blastocysts.

Besides overlapping DEGs with known bovine TF database, *de novo* motif calling enabled target-specific TF identification, revealing the most enriched sequence at DEG promoters. For example, *PABPC1,* also known as an embryonic poly(A)-binding protein, plays a key role in mRNA stabilization and translation initiation and is critical for preventing early embryonic arrest in human and mouse embryos [116]. Moreover, several TFs identified in promoters of sex-biased DEGs are critical to embryo development. For instance, *ETVs* (*ETV1, ETV5*), *ELK1*, *ELF1*, and *Sp1* are known to regulate the transcription of many genes during zygotic genomic activation (ZGA) [117–120]. The *KLF* family plays an important role in embryonic development and maintaining embryonic stem cell pluripotency [117, 120–122]. Interestingly, one common predicted motif from both sexes, *ELF4,* a key ZGA and epigenetic reprogramming regulator in pigs [123], was also identified as one of the female-biased DEGs in our analysis, suggesting that its expression also contributes to regulating its downstream genes. These results showed a more specific upstream regulation of sex-biased DEGs between male and female blastocysts.

### Sex-biased alternative splicing in early embryos

Alternative splicing plays a crucial role in transcriptome diversity by selectively including or excluding exons and introns from the same gene [124]. The resulting transcripts are often expressed in a developmental stage-specific manner and contribute to various biological processes, including gonadal differentiation [125], sex determination [126], and stress responses [127]. Importantly, AS helps in establishing the complex regulatory networks in mammals by generating multiple proteins from a single gene [128]. Sex-biased AS serves as an alternative mechanism for sexually dimorphic traits, complementing the sex-biased gene expression [129]. In Drosophila, sex-biased AS has been observed at the 0-2 hour stage, with alternative first exon usage as the most common type [130].

In this study, we identified more than 1,500 significant sex-biased AS events in bovine expanded blastocysts, with SE events being the most abundant. Specifically, male-biased SE events were more prevalent than those in females, which might due to that the increased activity of splicing regulators in female embryos could contribute to alternative exon exclusion. Analysis of protein domains affected by sex-biased SE events revealed distinct differences between male and female embryos. In male embryos, the most frequently skipped protein domains included FERM_N, which is involved in localizing proteins to the plasma membrane [131], HMG_box, which regulates DNA-dependent processes [132], and Nramp, which is associated with transmembrane transport [133]. In female embryos, the top skipped protein domains were DSPn, which plays a role in dephosphorylation [134], LRAT, which catalyzes vitamin A esterification [135], and Septin, which is essential for cytokinesis and other diverse cellular functions [136]. Moreover, functional enrichment analysis of all male-biased AS associated genes revealed significant correlations with mRNA metabolic process and mitochondrial functions, supporting the observation that energy metabolism was more active in male embryos. These findings highlight the potential role of AS in shaping sex-specific metabolic and developmental pathways during early embryogenesis and suggest that sex-biased AS events may lead to functional modifications in protein activity, potentially altering metabolic processes in female embryos.

### Sex-biased isoform expression in early embryos

By incorporating novel isoforms identified from long-read sequencing, we obtained a more comprehensive view of bovine embryonic isoform diversity, identifying more than 1,100 sex-biased isoforms that correspond to more than 1,000 genes. Considering both gene-level expression and isoform dynamics provided deeper insight into how different genes were finely tuned in a sex-specific manner. For example, *APOE*, which plays a crucial role in nerve system development during early development [137], exhibited male-biased expression at the gene level and also contained a male-biased isoform. In contrast, *KDM6A*, which is involved in cell differentiation and developmental gene regulation [138], showed female-biased expression at the gene level, but it did not contain any sex-biased isoforms.

Moreover, this study identified hundreds of genes exhibiting sex-biased expression only at the isoform level, without detectable differences at the gene level. These sex-biased isoforms may play key roles in sex-specific development, yet they would not have been detected by conventional differential gene expression analysis. One such example is *GOLM2*, a protein coding gene associated with Golgi apparatus [139], which contained a male-biased isoform but showed no sex biases at the gene level.

Examining the relationship between sex-biased isoforms and sex-biased AS events, we identified over 150 overlapped genes, suggesting that sex-biased isoforms contribute to shaping sex-biased AS patterns. Furthermore, functional enrichment of sex-biased isoforms associated genes revealed distinct differences between sexes. In female embryos, the most enriched functions were negative regulation of transcriptional activities and RNA biosynthetic process, while in male embryos, sex-biased isoform-associated genes were primarily involved in peptide metabolism, translation, and ATP metabolism. Overall, these findings provided novel insights into how sex-biased AS events, isoform expression, and gene-level expression collectively drive sex-specific early development.

### Conclusion

The faster development in male embryos compared to females has been observed across multiple species. However, the transcriptomic mechanisms driving these sex-specific differences in early embryo development remain incompletely understood. Through a comprehensive analysis of sex-biased DEGs, AS events, and DEIs in expanded blastocysts, we found that the faster cleaving of male embryos is driven by enhanced metabolic activity, particularly in amino acids and carbon metabolism. In contrast, female blastocysts exhibit slower cleavage, with an enrichment in female-specific developmental processes and pathways. We also identified several sex-specific transcription factors, RBPs, and splicing factors involved in DEG and AS regulation. Moreover, sex-biased AS events and DEIs play a critical role in fine-tuning sex-specific gene expression profiles, contributing to early developmental differences. Additionally, XCU and incomplete XCI appear to influence transcription activity in female embryos, potentially contributing to their developmental difference. This study provides new insights into how chromosomal composition and different levels of transcriptomic regulation contribute to sex differences in bovine early embryonic development, emphasizing the importance of considering sex as a factor in experimental design. Moreover, this study only focused on the expanded blastocyst stage. Future investigation examining earlier or later developmental stages is needed to provide a more comprehensive understanding of sex differences during early reprogramming and development.

## Declarations

### Ethics approval and consent to participate

Not applicable.

### Consent for publication

Not applicable.

### Availability of data and materials

The RNA-seq raw data have been deposited at NCBI Gene Expression Omnibus database GSE290265.The current study did not generate new code; all codes used for analysis in this study can be found according to corresponding references.

### Competing interests

The authors declare that they have no competing interests.

### Funding

This study was funded by the National Science Foundation through Grant # 2213824 to JED and Center for Vertebrate Genomics seed grant to SHC and JED.

### Authors’ contributions

Conceptualization: MS, GL, SHC and JED; Experimental work: MS, HA, and AL; Sample collection: MS, JZ, and LYL; Data analysis and interpretation: MS, GL, SHC, and JED; Funding: SHC and JED; Original draft writing: MS and GL; Revision of original draft: MS, GL, and JED; All authors read and approved the final version of the manuscript. Funding acquisition J.E.D. and the IISAGE Consortium.

## Acknowledgements

The author thanks all members of the IISAGE Consortium for the discussion of the results.

## Supplemental Materials

**Supplemental Figure S1. PCA of top 500 most variable genes in samples**.

**Supplemental Figure S2. PPI networks among male- and female-biased DEGs and non-DEGs.** Genes were ranked along the axis based on the degree of interactions.

**Supplemental Figure S3. Motif calling for female- and male-biased DEGs.** (A) Motifs at promoter regions of female-biased DEGs. (B) Motifs at promoter regions of male-biased DEGs. The motifs are clustered based on sequence similarity.

**Supplemental Figure S4. X:A ratio in bovine blastocysts**. The X:A ratio was recalculated in comparison of included or excluded the paralog genes in gene subgroups.

**Supplemental Figure S5. Distribution of Sex-biased DEGs on X-chromosome.** The left panel is sex-biased DEGs and the right panel is all anotated genes on X-chromosome. Gene density was calculated with 1Mb window. PAR: pseudoautosomal region.

**Supplemental Figure S6. Binding potential of Splicing factors FMR1 and HNRNPH2 on sex-biased SE events.** The green block indicated the alternative exons in the sex-biased SE events, with the negative numbers marking the upstream region of the alternative exons and the positive numbers marking the downstream region of the alternative exons. The red line indicated the binding potential for female-biased SE events, and the blue line indicated the binding potential for male-biased SE events. The significance of the splicing factors binding was shown on the right axis.

**Supplemental Figure S7. Features of long-read isoforms.** (A) Illustrations of different categories for long-read isoforms, including full splice match FSM), incomplete splice match (ISM), novel in catalog (NIC), novel not in catalog (NNC), genic Intron, and genic genomic. (B) Number of long-read isoforms in each category. (C) Number of isoforms per gene based on long-read data. (D) The percent of transcripts with annotation supported isoforms (Annotated), canonical junctions (Canonical), and splicing junctions with short-read coverage (Coverage SJ) for the top 4 isoforms categories (NNC, FSM, ISM, and NIC).

**Supplemental Figure S8. Expression pattern at the gene level and the associated isoforms.** (A) Gene level and isoform level expression for *APOE*. (B) Gene level and isoform level expression for *KDM6A*. (C) Gene level and isoform level expression for *GOLM2*. *: padj < 0.05; ns: non-significant.

**Supplemental Figure S9. The sex-biased exon usage in *DENND1B* and *DIS3L2*.** (A) Significant female-biased exon usage (Exon 7) in *DENND1B* on negative strand. (B) Significant female-biased exon usage (Exon 13) in *DIS3L2* on positive strand. The top line plot showed the exon usage information between sexes, and the purple block below indicated the exons with significantly differential usages between sexes.

**Supplemental Figure S10. PCR validation of SE events.** The SE events were validated in *CEP350*, *ZMYND11*, and *WWC2*. The left three panels were the gel image of the PCR products, while the right panels were the example to show the skipped exon highlighted in green. F1, F2, and F3 are female embryos; while M1, M2, and M3 are male embryos.

**Supplemental Figure S11. AS validation.** This bar plot compared the inclusion level between the PCR validation and RNA-seq data. AS events between sexes were compared via the inclusion levels, which is the ratio of the expression level of the transcripts with the skipped exon to the expression level of the total he transcripts. The expression level of PCR validation was calculated based on the brightness of the band with Image-J.

**Supplemental Table S1. Primer information for embryo determination**

**Supplemental Table S2. Primer information for sex-biased DEG validation**

**Supplemental Table S3. Primer information for alternative splicing validation**

**Supplemental Table S4. Sex ratio of bovine IVF embryos and summary statistics of RNA-seq data**

**Supplemental Table S5. Differentially expressed genes and functional enrichment**

**Supplemental Table S6. Overlapped genes between sex-biased DEGs and AS genes and functional enrichment**

**Supplemental Table S7. Sex-biased alternative splicing events**

**Supplemental Table S8. Protein domains affected by sex-biased SE events and functional enrichment**

**Supplemental Table S9. Sex-biased isoforms and functional enrichment**

**Supplemental Table S10. Overlapping genes among sex-biased DEGs, differential alternative splicing genes, and differential isoform-related genes.**

**Supplemental Table S11. Differential exon usage between sexes**.

## Notes

### Competing Interest Statement

The authors have declared no competing interest.

