## Supplementary figures and images for "Sex-biased Transcriptome in *in vitro* Produced Bovine Early Embryos"

### Figure S1

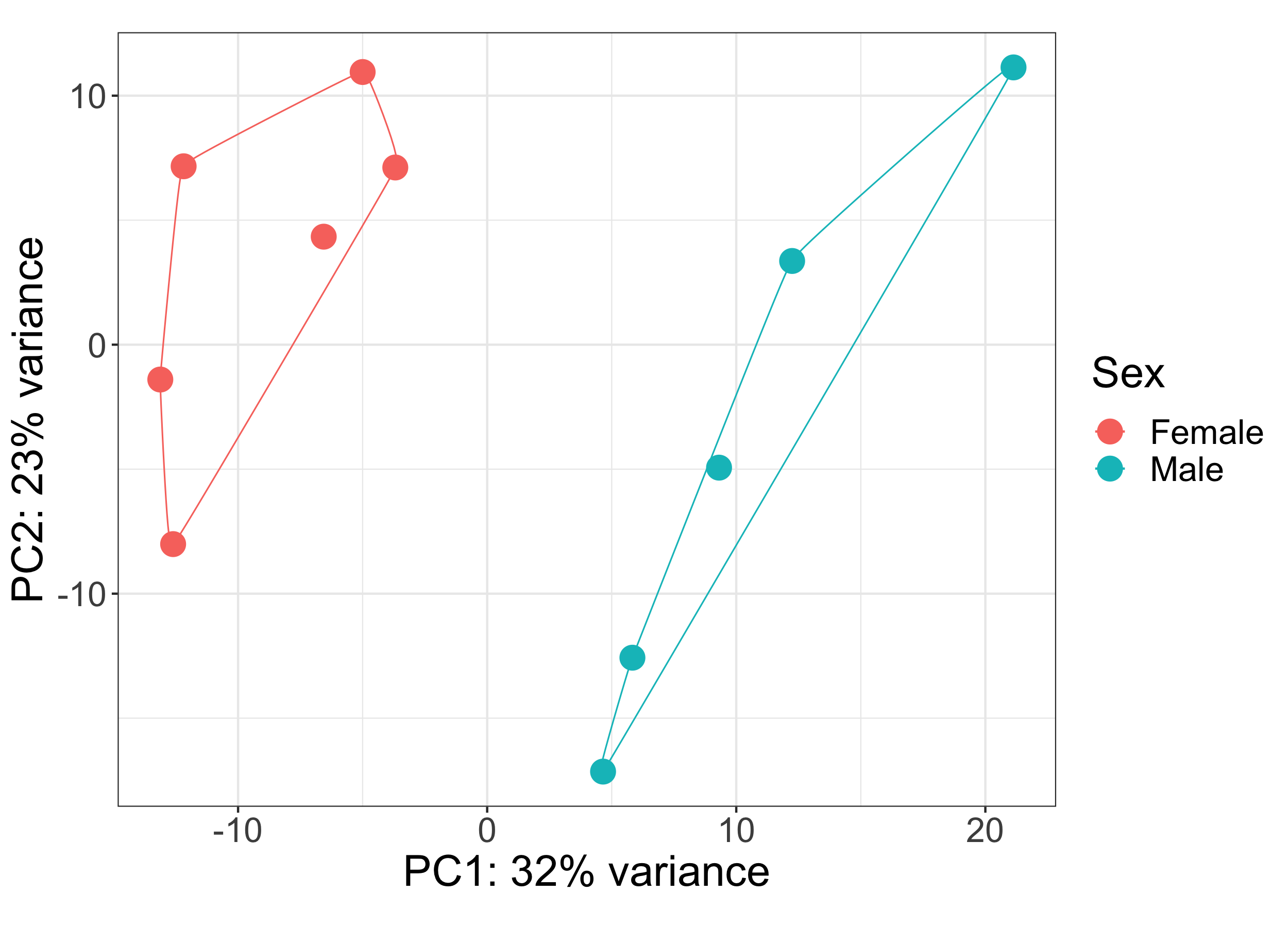

### Figure S2

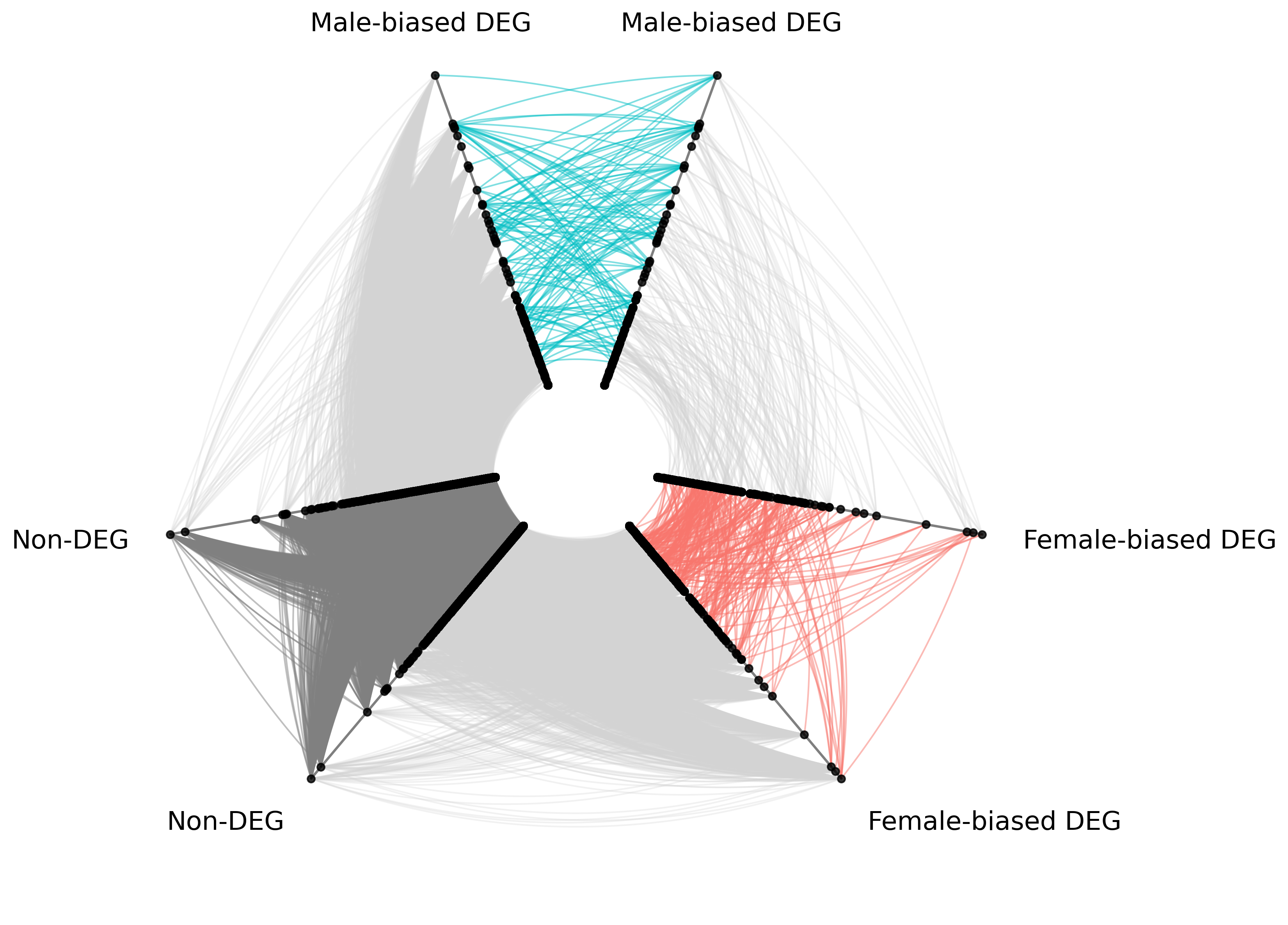

### Figure S3

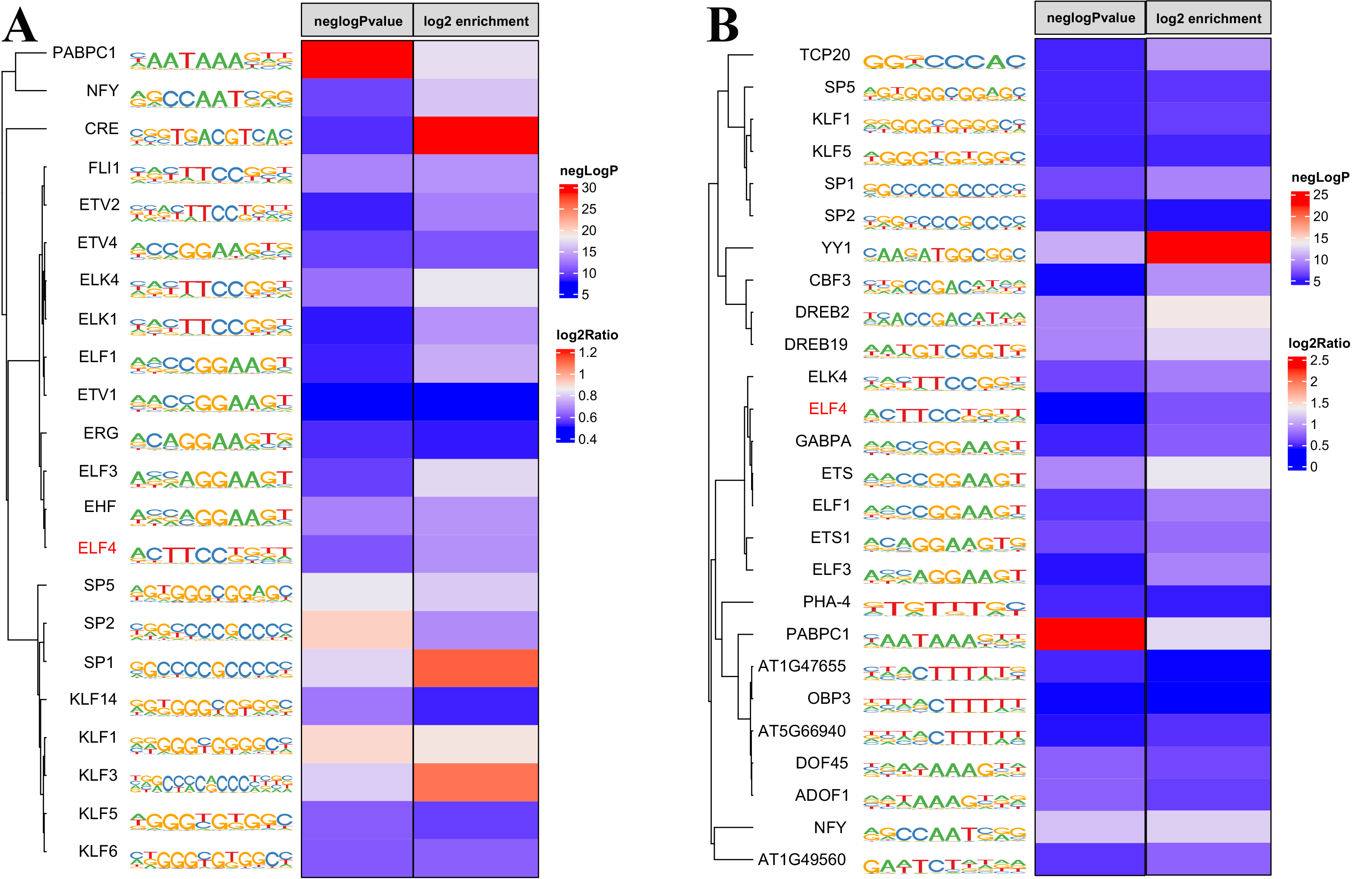

### Figure S4

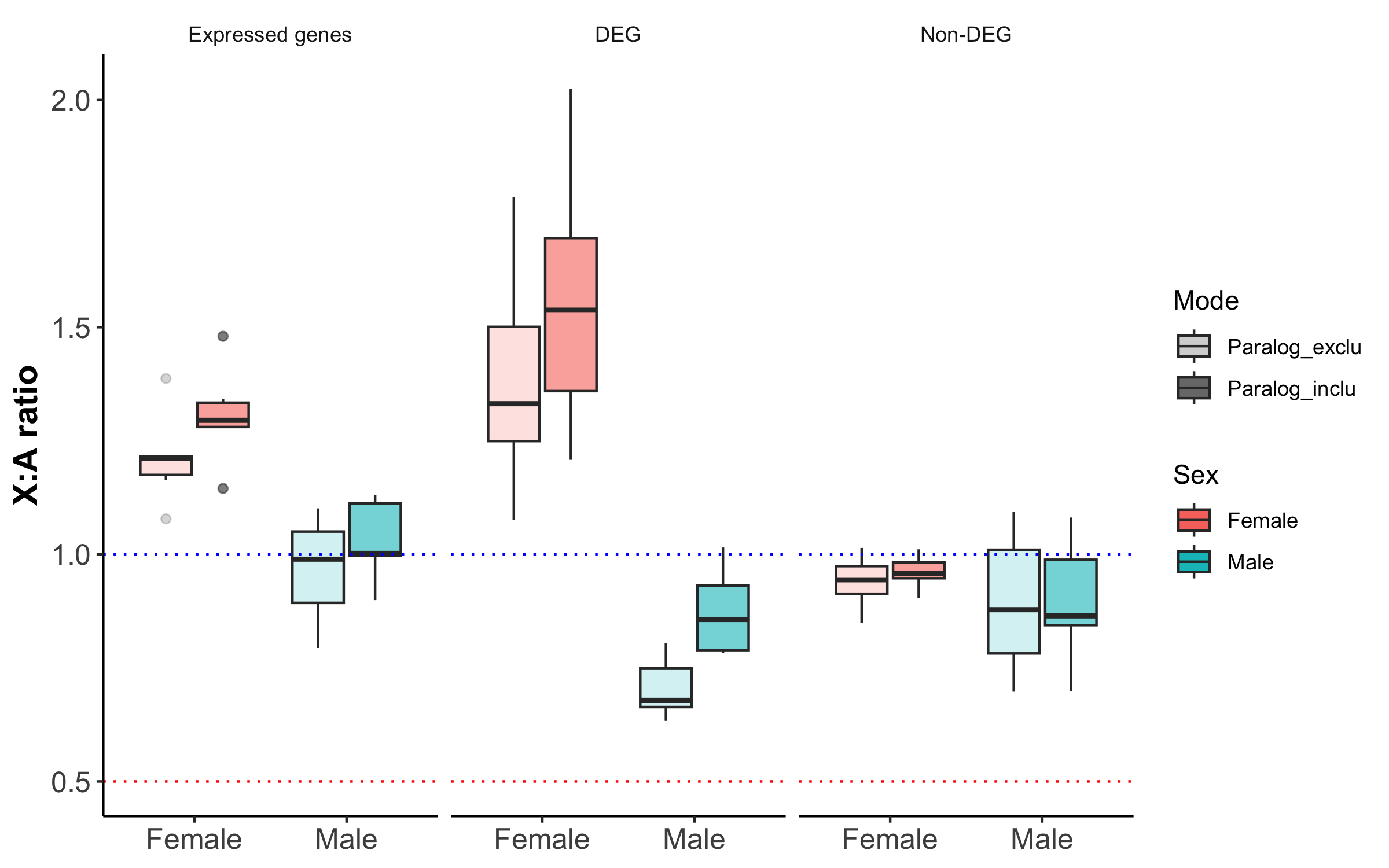

### Figure S5

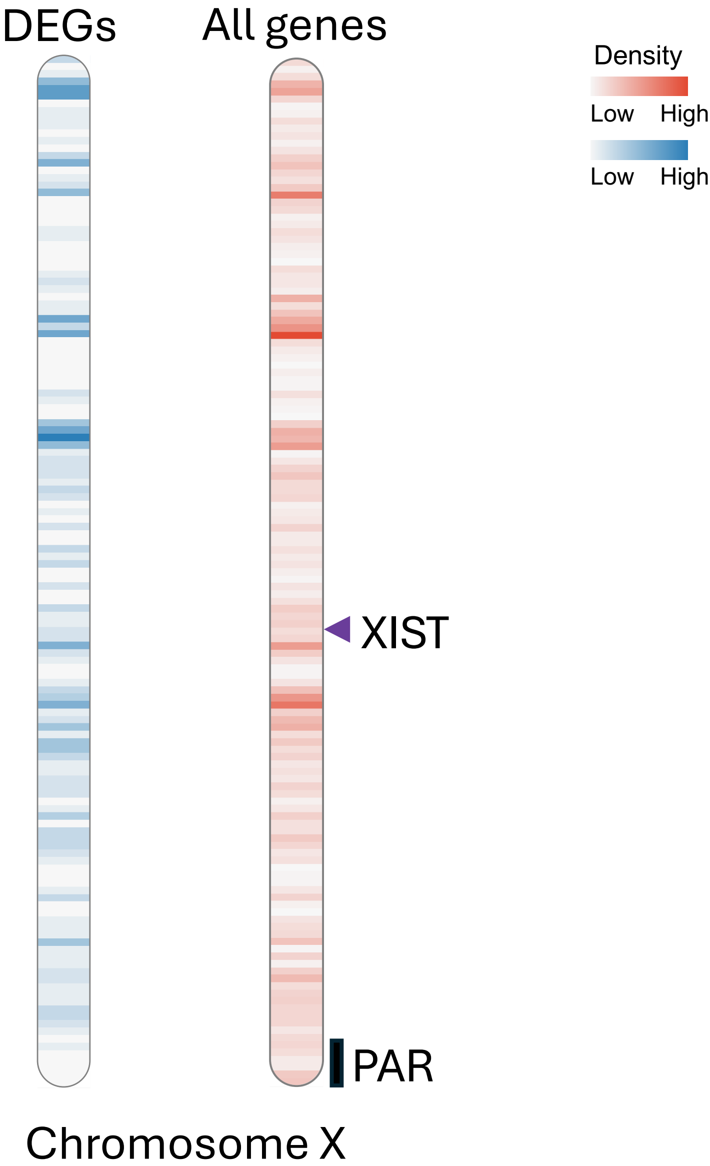

### Figure S6

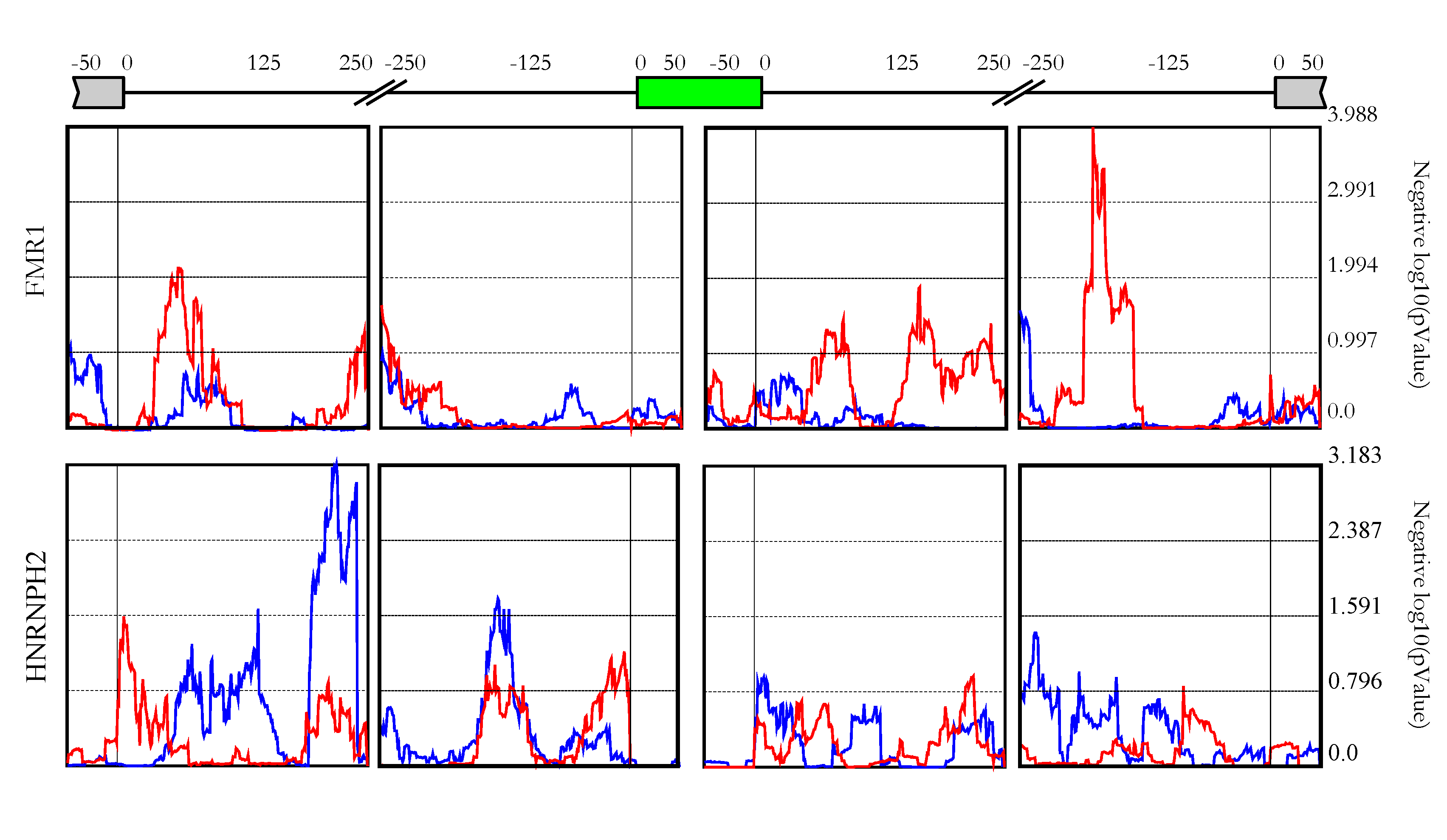

### Figure S7

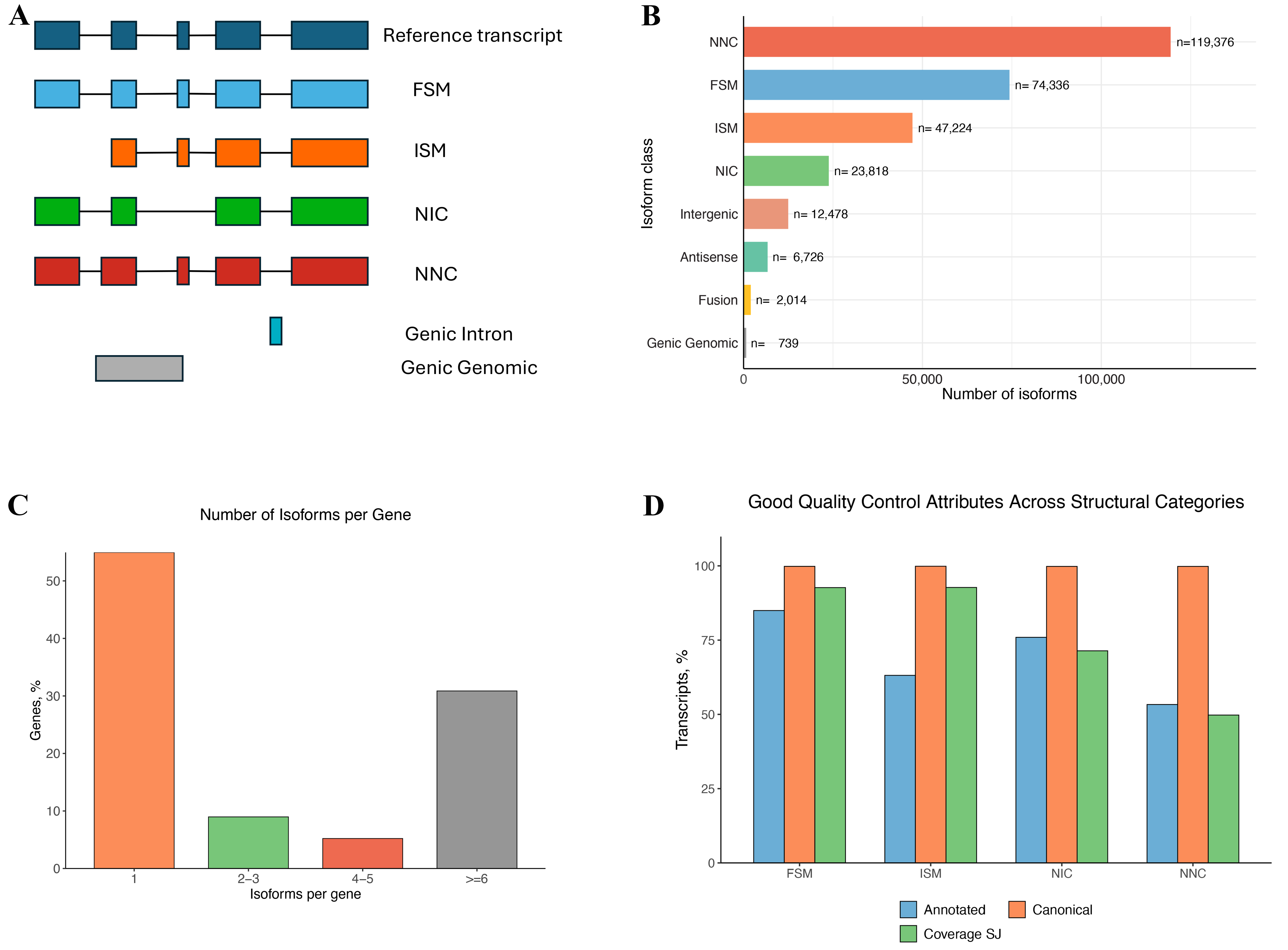

### Figure S8

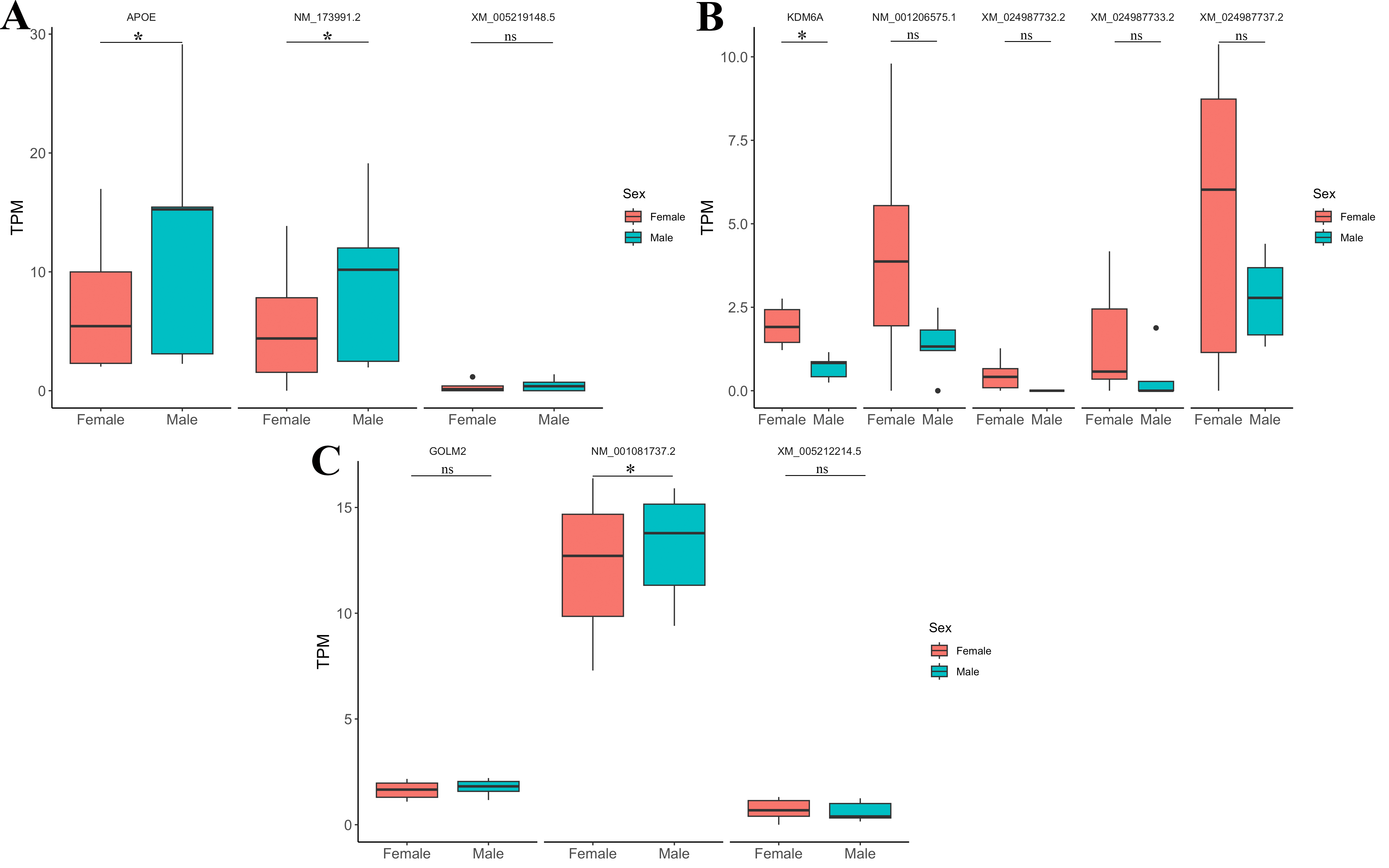

### Figure S9

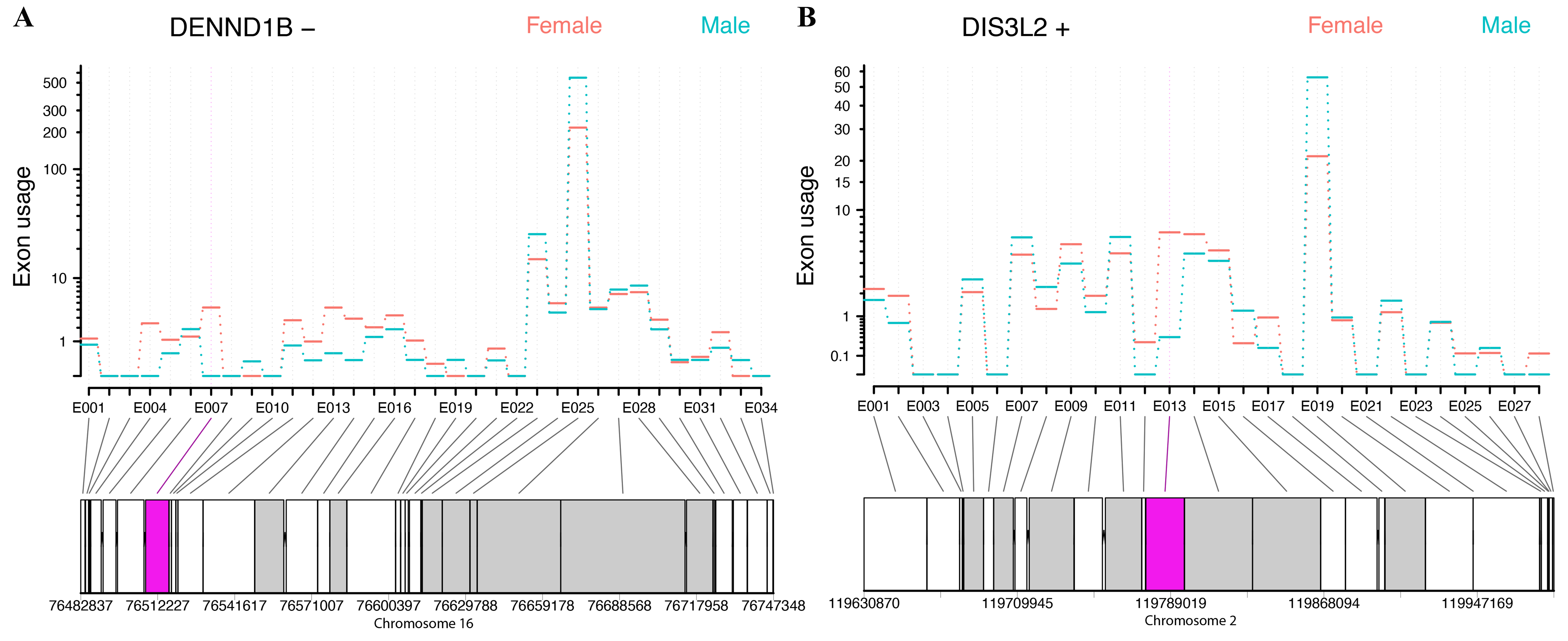

### Figure S10

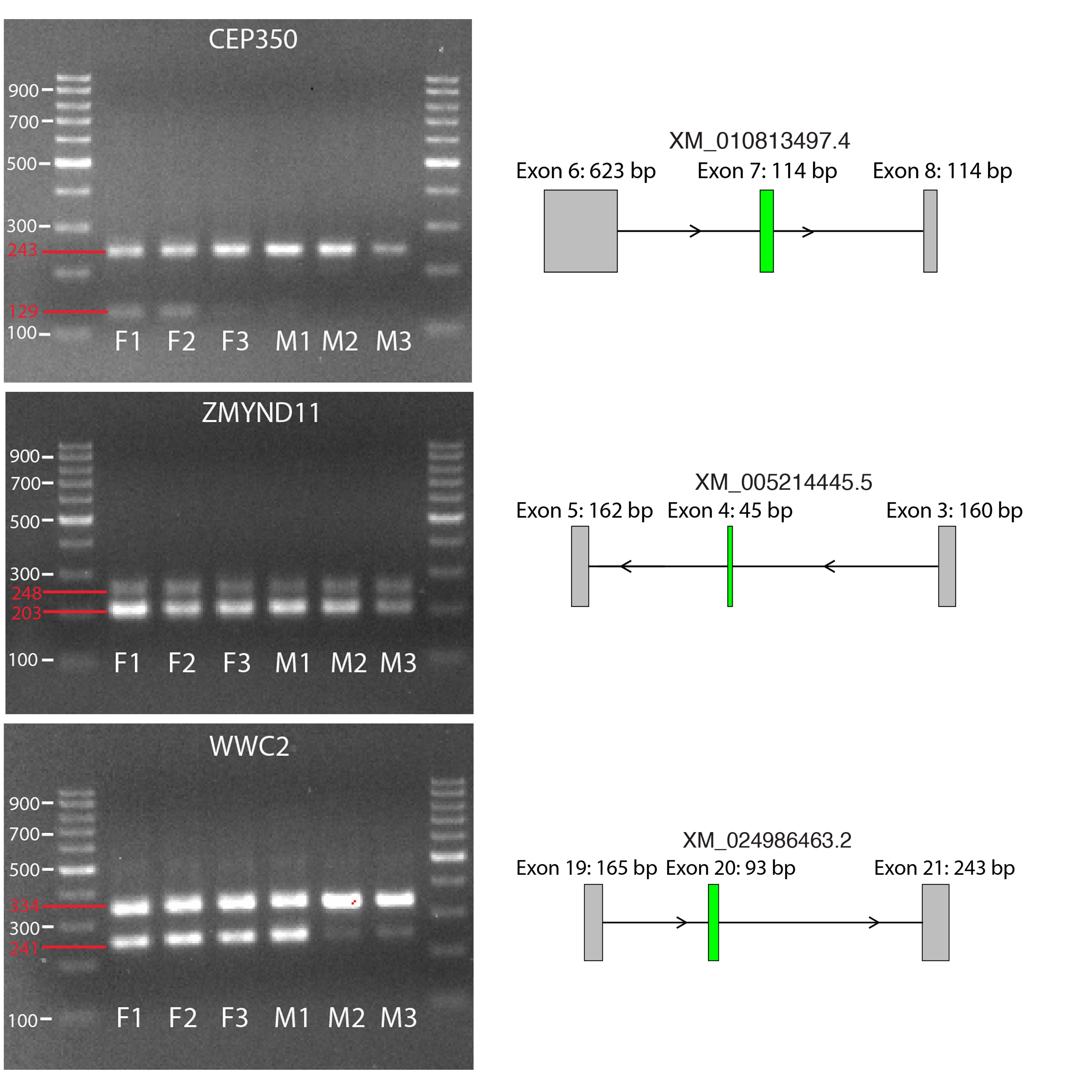

### Figure S11

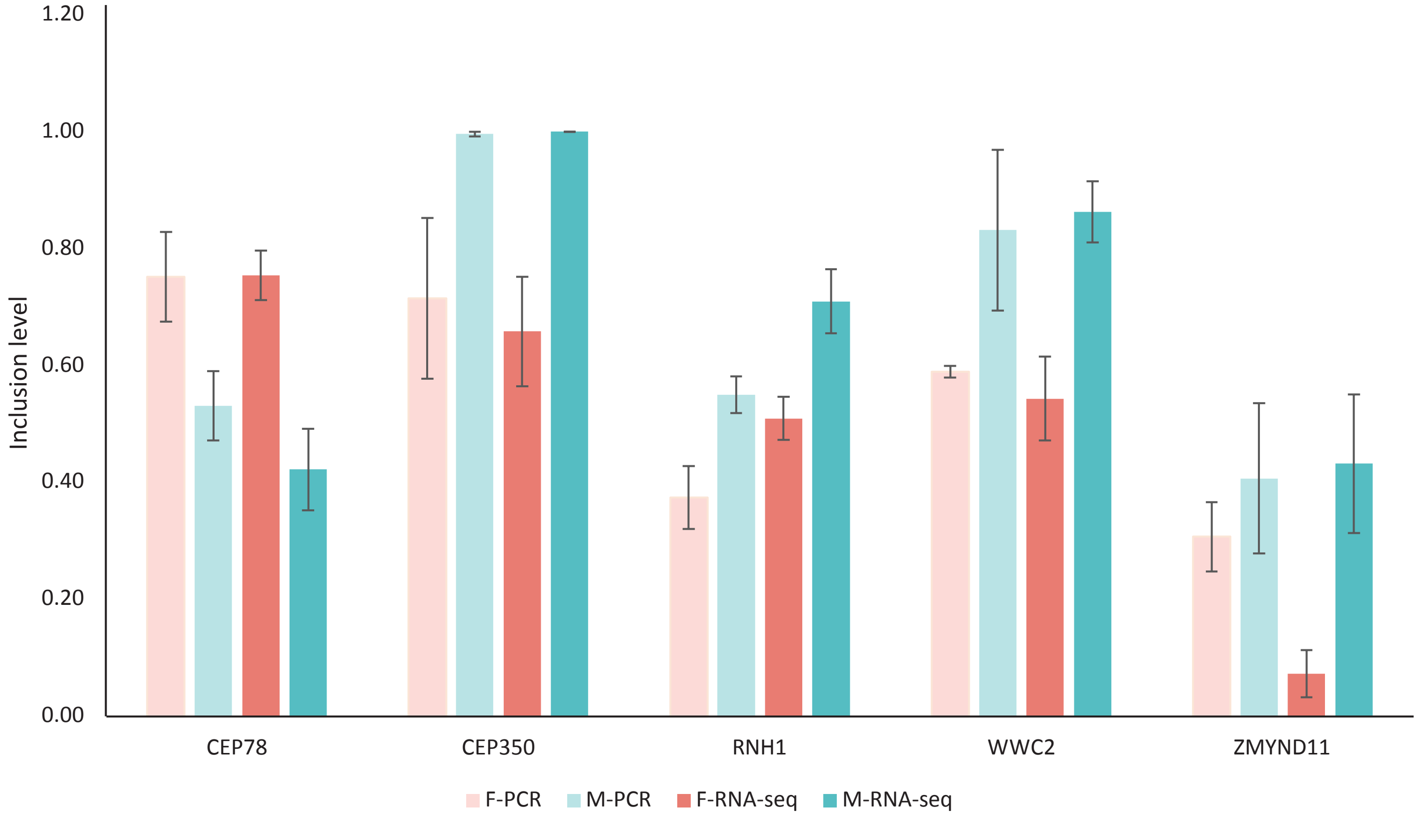
